# Bayesian model selection favors parametric over categorical fMRI subsequent memory models in young and older adults

**DOI:** 10.1101/2020.07.27.220871

**Authors:** Joram Soch, Anni Richter, Hartmut Schütze, Jasmin M. Kizilirmak, Anne Assmann, Lea Knopf, Matthias Raschick, Annika Schult, Anne Maass, Gabriel Ziegler, Alan Richardson-Klavehn, Emrah Düzel, Björn H. Schott

**Affiliations:** German Center for Neurodegenerative Diseases (DZNE), Göttingen, Germany; Bernstein Center for Computational Neuroscience (BCCN), Berlin, Germany; Leibniz Institute for Neurobiology (LIN), Magdeburg, Germany; German Center for Neurodegenerative Diseases (DZNE), Magdeburg, Germany; Otto von Guericke University, Medical Faculty, Magdeburg, Germany; Independent scholar, Berlin, Germany; Center for Behavioral Brain Sciences (CBBS), Magdeburg, Germany; Department of Psychiatry and Psychotherapy, University Medical Center Göttingen, Göttingen, Germany

**Keywords:** subsequent memory effect, Bayesian model selection, episodic memory, parametric fMRI, aging

## Abstract

Subsequent memory paradigms allow to identify neural correlates of successful encoding by separating brain responses as a function of memory performance during later retrieval. In functional magnetic resonance imaging (fMRI), the paradigm typically elicits activations of medial temporal lobe, prefrontal and parietal cortical structures in young, healthy participants. This categorical approach is, however, limited by insufficient memory performance in older and particularly memory-impaired individuals. A parametric modulation of encoding-related activations with memory confidence could overcome this limitation. Here, we applied cross-validated Bayesian model selection (cvBMS) for first-level fMRI models to a visual subsequent memory paradigm in young (18-35 years) and older (51-80 years) adults. Nested cvBMS revealed that parametric models, especially with non-linear transformations of memory confidence ratings, outperformed categorical models in explaining the fMRI signal variance during encoding. We thereby provide a framework for improving the modeling of encoding-related activations and for applying subsequent memory paradigms to memory-impaired individuals.

## 1. Introduction

The subsequent memory paradigm, the comparison of encoding-related brain responses to stimuli as a function of their later remembering or forgetting, is a widely used approach in neuroimaging research of human explicit and particularly, episodic, memory. The neural signatures that differentiate subsequently remembered from subsequently forgotten stimuli are commonly referred to as the DM effect (difference [due to later] memory; Paller et al., 1987). First employed in human event-related potential studies (Paller et al., 1987), the DM approach has been established as a key paradigm in event-related functional magnetic resonance imaging (fMRI) since the publication of two landmark studies over two decades ago (Brewer, 1998; Wagner et al., 1998). In a typical fMRI study of successful memory formation, the DM effect is experimentally evoked by presenting a subject with novel information (*encoding*) and assessing encoding success in a subsequent memory test (*retrieval*). During encoding, subsequently remembered stimuli elicit increased brain responses in the hippocampus and adjacent medial temporal lobe (MTL) structures as well as in prefrontal and occipito-parietal brain structures when compared to subsequently forgotten items, and these findings have been robustly replicated in numerous studies (for a meta-analysis, see Kim, 2011). Over the past two decades, variations of the subsequent memory paradigm have been adapted to a variety of questions in cognitive memory research, like the common and distinct processes of implicit and explicit memory (Schott et al., 2006; Turk-Browne et al., 2006), the dissociation of encoding processes related to later recollection and familiarity (Davachi et al., 2003; Henson et al., 1999), or the influence of different study tasks on neural correlates of encoding (Fletcher et al., 2003; Otten and Rugg, 2001). While most of those studies have been conducted in young, healthy adults, the DM paradigm has also been successfully applied to older adults (Düzel et al., 2011; for a review, see Maillet and Rajah, 2014) or to clinical populations, such as patients with temporal lobe epilepsy (Richardson et al., 2003; Towgood et al., 2015) or schizophrenia (Bodnar et al., 2012; Zierhut et al., 2010).

Episodic memory performance declines during aging, and previous studies suggest that age-related changes in encoding-related brain activity differ between individuals with rather preserved memory function (“successful aging”) and subjects with relevant age-related memory decline (Düzel et al., 2011; Maillet and Rajah, 2014). Therefore, the fMRI DM effect might be useful in assessing a potential neural underpinning of individual differences in age-related alterations of the MTL memory system. However, when applying the subsequent memory paradigm to older participants with poor memory performance, a limitation arises from the fact that older subjects and particularly those with memory dysfunction often remember an insufficient number of stimuli to allow for a meaningful comparison of later remembered and later forgotten items, whereas young healthy participants might conversely achieve ceiling performance in easier memory tasks, rendering it equally difficult to infer on subsequent memory effects.

In subsequent memory experiments, encoding success can be assessed via different retrieval tasks, which are commonly based on either recall (free or cued) or recognition. In recognition paradigms, previously presented (*old*) and previously unseen stimuli (*new*) are shown in random order, and subjects are asked whether they have seen an item during encoding or not. Some recognition memory tests do not merely rely on binary responses, but instead require subjects to provide a recognition confidence rating on a Likert scale (Likert, 1932) by, for example, judging items on a five-point scale from “definitely old” via “probably old”, “uncertain”, and “probably new” to “definitely new”. This approach has been used to infer on neural correlates of recognition, that is, familiarity (differentiation of old/new without reporting additional details from the encoding episode) and recollection (recognition memory accompanied by remembering of contextual details of the encoding episode) (Düzel et al., 2011; Schoemaker et al., 2014).

When assuming that the majority of older and even memory-impaired individuals exhibit at least some variability in responding on such a scale (e.g., from “definitely new” to “uncertain”), one could model the subsequent memory effect parametrically. While categorical models employ variables that can only assume nominal values (e.g., remembered vs. not remembered), parametric models employ variables with a wider ranges of values, (e.g., the degree of confidence whether an item is remembered). The usefulness of parametric models is two-fold: Firstly, participants are no longer required to make a definite decision when they are actually uncertain. Secondly, the parametric models are inherently less complex, as they can incorporate multiple responses in a single regressor. This lower complexity, however, relies on the assumption that the relationship between the parametric regressor and the measured response is itself parametric in nature (Bogler et al., 2013; Soch et al., 2016, Fig. 3B; Soch et al., 2020, Fig. 8C). Several previous studies have provided promising evidence for the applicability of parametric analyses to fMRI-based DM effects (Dennis et al., 2008; Fernández et al., 1998; Kim and Cabeza, 2007; Richter et al., 2017), but it should be noted non-linear parametric modulations may be superior to simple linear parametric regressors (Daselaar et al., 2006).

To date, the use of parametric approaches in analyzing subsequent memory fMRI data has not undergone an objective validation. A parametric analysis would be based on the assumption that the BOLD signal in memory-related brain regions varies quantitatively rather than qualitatively with the strength of the encoding signal. It may therefore potentially be suboptimal when considering multi-process models of explicit memory, such as the dual-process signal detection model of recollection and familiarity (Yonelinas, 1994; Yonelinas et al., 2010). On the other hand, parametric models could outperform categorical models due to their lower complexity. Furthermore, when using confidence scales that allow for uncertain responses or guesses, parametric models might also be employed in memory-impaired subjects whose behavioral performance does not allow for meaningful categorical modeling of the DM effect.

Here, we used an objective model selection approach to explore the applicability of parametric compared to categorical models of the fMRI subsequent memory effect, using a visual memory encoding task with a five-point confidence rating during a recognition memory test that has previously been employed to assess neural correlates of successful aging (Düzel et al., 2011, Fig. 1). Subject-wise general linear models (GLMs; Friston et al., 1994) of individual fMRI datasets were treated as generative models of neural information processing, and the selection between the different GLMs was afforded by voxel-wise cross-validated Bayesian model selection (cvBMS; Soch et al., 2016). This approach results in an estimated frequency for each model that informs us how often this model provides the optimal explanation of the observed data. We hypothesized that models including a differentiation of subsequently remembered and subsequently forgotten items would outperform models that did not account for memory performance and that among these models, parametric models would be superior to categorical models of successful encoding.

**Figure 1.**
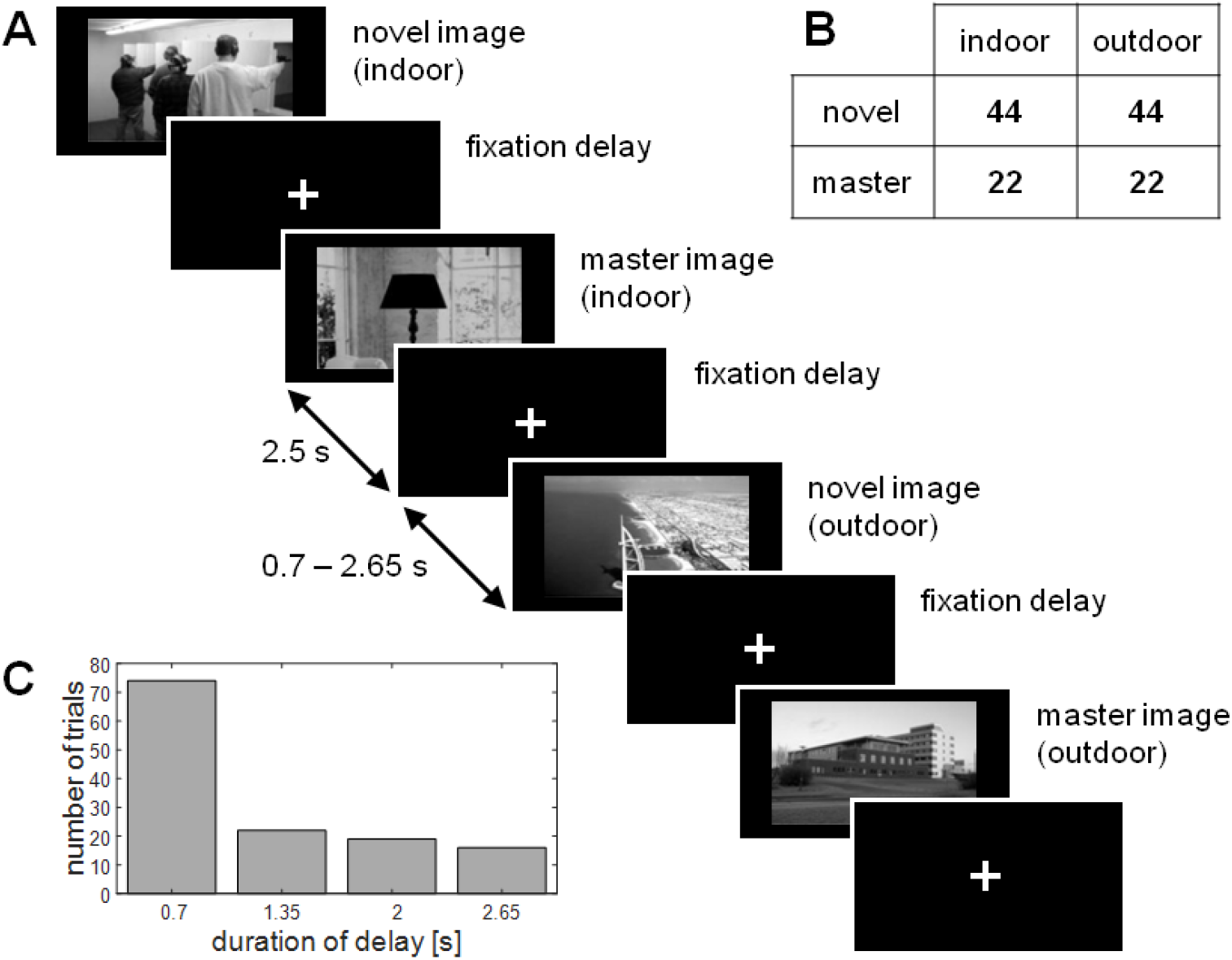
Experimental design and stimulus timing during encoding. **(A)** Exemplary sequence of trials, each trial consisting of either a previously unseen novel image or a pre-familiarized master image showing either an indoor or an outdoor scene. Each stimulus was shown for 2.5 s and followed by a variable inter-stimulus-interval (ISI) between 0.7 and 2.65 s. **(B)** Number of trials in the four experimental conditions. There were equally many indoor and outdoor scences and twice as many novel images as repetitions of the two previously familiarized master images. **(C)** Distribution of ISIs in the encoding session. ISIs were pseudo-exponentially distributed with shorter intervals occurring more often than longer ones.

## 2. Methods

### 2.1. Participants

A total of 279 volunteers participated in the study (117 young, 162 older; see Supplementary Methods for sample size estimation). Data from 20 participants had to be excluded from analysis due to history of psychiatric conditions (five cases), incidental findings in structural MRI scans (eight cases), technical difficulties during recording of behavioral responses and/or MRI of the memory experiment (four cases), nausea during scanning, insufficiently corrected vision, or artifacts in the MR images (one case each). The resulting study cohort consisted of a total of 259 neurologically and psychiatrically healthy adults, including 106 young (47 male, 59 female, age range 18-35, mean age 24.12 ± 4.002 years) and 153 older (59 male, 94 female, age range 51-80, mean age 64.04 ± 6.735 years) participants. The study was approved by the Ethics Committee of the Otto von Guericke University Magdeburg, Faculty of Medicine, and written informed consent was obtained from all participants in accordance with the Declaration of Helsinki (World Medical Association, 2013).

### 2.2. Experimental paradigm

During the fMRI experiment, participants performed a visual memory encoding paradigm with an indoor/outdoor judgment as the incidental encoding task (see Figure 1A). Compared to earlier publications of this paradigm (Assmann et al., 2020; Barman et al., 2014; Düzel et al., 2011; Schott et al., 2014), the trial timings had been adapted as part of the DELCODE protocol (Bainbridge et al., 2019; Düzel et al., 2019). Subjects viewed a series of photographs showing either an indoor or an outdoor scene, which were either novel to the participant at the time of presentation (44 indoor and 44 outdoor scenes) or were repetitions of two pre-familiarized “master” images (i.e. one indoor and one outdoor scene shown to the participants before the start of the actual experiment; see Figure 1B). Irrespective of novelty, subjects were requested to categorize images as “indoor” or “outdoor” via button press. Each picture was presented for 2.5 s, followed by a variable delay between 0.70 s and 2.65 s (see Figure 1C), with stimulus intervals and order optimized for an efficient estimation of the trial-specific BOLD responses (Düzel et al., 2011; Hinrichs et al., 2000).

Approximately 70 minutes (70.23 ± 3.77 min) after the start of the fMRI session, subjects performed a recognition memory test outside the scanner, in which they were presented with photographs that had either been shown during the fMRI experiment or were novel to the participant at the time of presentation. Among the 134 pictures presented to each subject during retrieval, 88 were previously seen “target” images (44 indoor and 44 outdoor scenes), 44 were previously unseen “distractor” images (22 indoor and 22 outdoor scenes), and 2 were the previously seen pre-familiarized “master” images (1 indoor and 1 outdoor scene).

Subjects were requested to provide a recognition memory confidence rating using a five-point Likert scale with the following levels:

1. I am sure that this picture is new (*definitely new*).
2. I think that this picture is new (*probably new*).
3. I cannot decide if the picture is new or old (*unsure*).
4. I think I have seen this picture before (*probably old*).
5. I am sure that I have seen this picture before (*definitely old*).

The responses during this retrieval session were provided verbally by the participant and recorded via button press by an experimenter. These data were used to model the DM effect (see Section 3).

### 2.3. fMRI data acquisition

Structural and functional MRI data were acquired on two Siemens 3T MR tomographs, (Siemens Verio; 58 young, 83 older; Siemens Skyra: 48 young, 70 older), following the exact same protocol used in the DELCODE study (Düzel et al., 2019; Jessen et al., 2018).^1^

For structural MRI (sMRI), a T1-weighted MPRAGE image (TR = 2.5 s, TE = 4.37 ms, flip-α = 7°; 192 slices, 256 × 256 in-plane resolution, voxel size = 1 × 1 × 1 mm) was acquired for later co-registration. Phase and magnitude fieldmap images were acquired to improve spatial artifact correction (*unwarping*, see below).

For functional MRI (fMRI), 206 T2*-weighted echo-planar images (TR = 2.58 s, TE = 30 ms, flip-α = 80°; 47 slices, 64 × 64 in-plane resolution, voxel size = 3.5 × 3.5 × 3.5 mm) were acquired in interleaved-ascending slice order (1, 3, …, 47, 2, 4, …, 46). The total scanning time during the task-based fMRI session was 531.48 s. The complete study protocol also included a resting-state fMRI (rs-fMRI) session comprising 180 scans and using the same scanning parameters as in task-based fMRI (Teipel et al., 2018) as well as additional structural imaging (FLAIR, FLASH, susceptibility-weighted imaging; see e.g. (Betts et al., 2019), which are not subject of the analyses reported here.

### 2.4. fMRI data preprocessing

Data preprocessing and analysis were performed using Statistical Parametric Mapping (SPM12; Wellcome Trust Center for Neuroimaging, University College London, London, UK). First, functional scans (EPIs) were corrected for acquisition time delay (*slice timing*), followed by a correction for head motion (*realignment*) and magnetic field inhomogeneities (*unwarping*), using voxel-displacement maps (VDMs) derived from the fieldmaps. Then, the MPRAGE image was spatially co-registered to the mean unwarped image and segmented into six tissue types, using the unified segmentation and normalization algorithm implemented in SPM12. The resulting forward deformation parameters were used to normalize unwarped EPIs into a standard stereotactic reference frame (Montreal Neurological Institute, MNI) using a target voxel size of 3×3×3 mm. Finally, normalized images were spatially smoothed using an isotropic Gaussian kernel with full width at half maximum (FWHM) of 6 mm

### 2.5. Bayesian model selection

After preprocessing, fMRI data were analyzed using a set of first-level GLMs that provided the model space for the following model selection procedure (see Section 3). Model inference was performed via cvBMS (Soch et al., 2016) implemented in the SPM toolbox for model assessment, comparison and selection (MACS; Soch and Allefeld, 2018). Model inference either addressed individual GLMs, applied to voxel-wise cross-validated log model evidences (cvLME), or families of GLMs, applied to voxel-wise log family evidences (LFE) calculated from cvLMEs.

At the second level, cvBMS uses random-effects Bayesian model selection (RFX BMS), a hierarchical Bayesian population proportion model, the results of which characterize how prevalent each model (or model family) is in the population investigated. A proportion resulting from cvBMS (e.g. the likeliest frequency, LF) – can be interpreted as (i) the frequency of the population “using” a particular model or as (ii) the probability that a particular model is the generating model of the data of a given single subject. Consequently, the model with the maximum LF outperforms all other models in terms of relative frequency and may be regarded as the selected model in a cvBMS analysis. For each analysis reported in the results section, we show LF-based selected-model maps (SMM) scaled between 0 and 1, which display the most prevalent model in each voxel (Soch et al., 2016).

Note that, when interpreting SMMs, they should not be confused with statistical parametric maps (SPMs) conventionally reported as fMRI results. Whereas voxels on a thresholded SPM usually indicate that there is a significant activity difference in these voxels, a voxel appearing on an SMM only indicates that the respective model (or model family) performs better in explaining this voxel’s time course, relative to some other model. Consequently, the more brain regions appear on an SMM, the more evidence at the whole-brain level that this model is the optimal description of the neural processing underlying the cognitive task.

### 2.6. Replication in independent cohort

The paradigm employed in the present study had previously been used in another cohort of 117 young subjects (Assmann et al., 2020; see Supplementary Online Material, Table S1 and Figure S1). In the present study, we used those previously acquired datasets as an independent cohort for replication of the results obtained from the young subjects. All core findings could be replicated in that cohort, despite a small difference in trial timings. Results from the model selection analyses performed in the replication cohort are displayed in Supplementary Figures S5-S10, which are designed analogously to Figures S3 and 3-7 in the main manuscript.

## 3. Analysis

Preprocessed fMRI data were analyzed using first-level voxel-wise GLMs that were then submitted to cvBMS. In total, the model space consisted of 19 models (see Table 1 and Supplementary Figure S2), varying in their modeled event duration, categorization of trials and modeling of the subsequent memory effect.

**Table 1.**
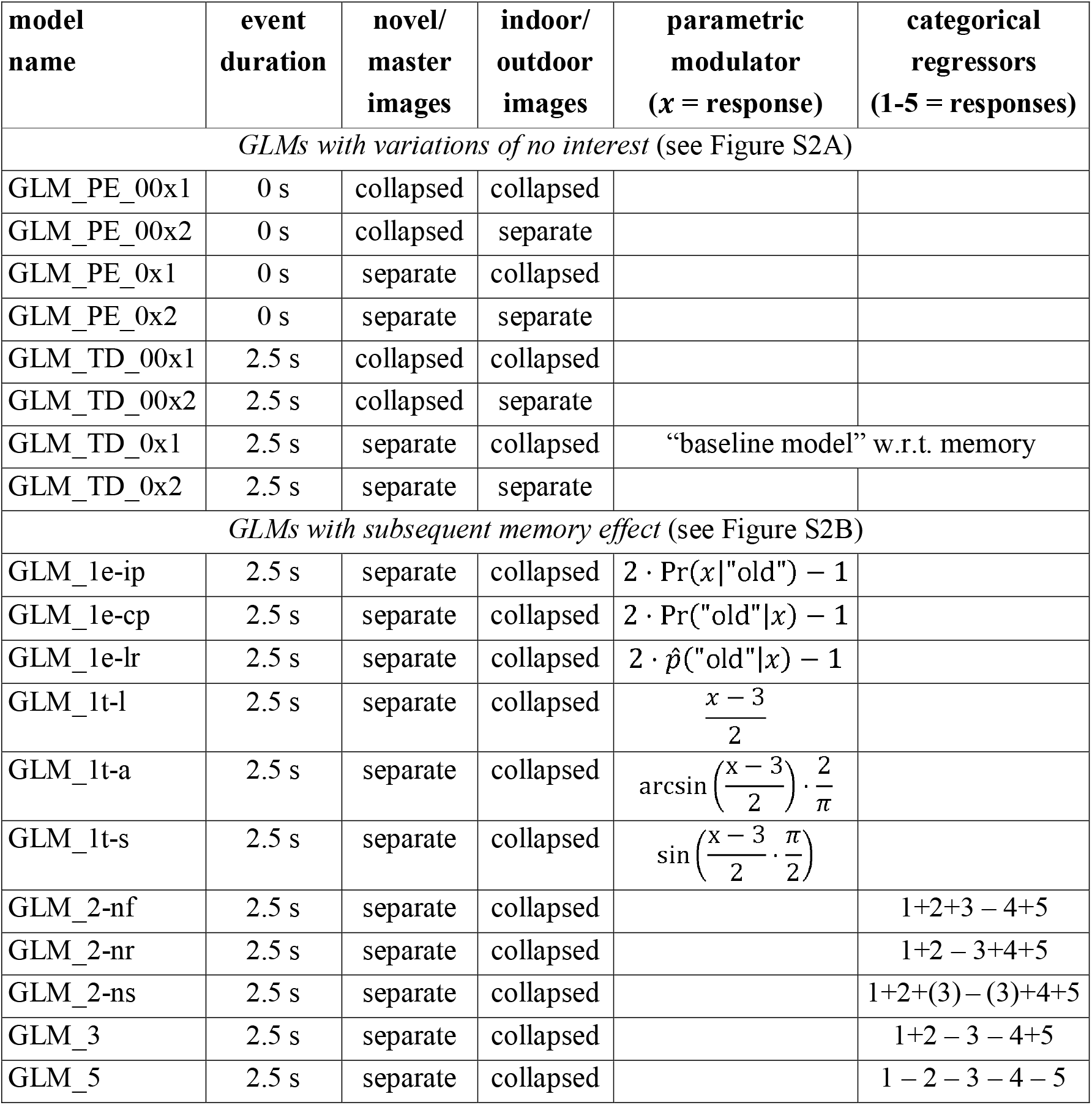
Model space for GLM-based fMRI analyses. 8 models without memory effects varying model features of no interest (top) and 11 models varying by the way how memory effects are modelled (bottom). Model names correspond to those in Supplementary Figure S2 and descriptions of memory regressors are given in Section 3.

### 3.1. The baseline model and variations of no interest

We began our GLM-based analysis by specifying the most straightforward model, in line with standard fMRI analysis conventions and most suitable for inferring novelty-related effects. This baseline model (marked as red in Figure S2A) included two onset regressors, one for novel images at the time of presentation (*novelty* regressor) and one for the two pre-familiarized images (*master* regressor). Both regressors were created as stimulus functions with an event duration of 2.5 s, convolved with the canonical hemodynamic response function, as implemented in SPM. Additionally, the model included the six rigid-body movement regressors obtained from realignment and a constant regressor representing the implicit baseline.

The baseline GLM was then varied along three modeling dimensions of no interest (see Table 1 and Figure S2A) that served for control and validation purposes (see Section 4.1):

- Stimulus-related brain responses can be either modeled according to the actual trial duration (TD) of 2.5 s (family *GLMs_TD* including the baseline GLM) or trials can be modeled as point events (PE) with a duration of 0 s, i.e. as delta functions (family *GLMs_PE*), resulting in shorter BOLD responses in the HRF-convolved regressors.
- Novel and master images can be either separated into two regressors (family *GLMs_0* including the baseline GLM) or events can be collapsed across these two conditions, yielding one single regressor (family *GLMs_00*).
- Indoor and outdoor scenes can be either collected into one regressor (family *GLMs_x1* including the baseline GLM) or events can be grouped into indoor and outdoor stimuli, yielding two regressors per condition (family *GLMs x2*).

Applying these three variations to the baseline GLM results in a model space of 2^3^ = 8 models (see Table 1), which allows to infer on the optimal event duration (0 s vs. 2.5 s), the novelty effect (novelty/master separated vs. collapsed) and the indoor/outdoor effect (indoor/outdoor separated vs. collapsed) by appropriate comparison of the model families.

Related to memory, the baseline GLM allows inferring on a novelty effect by contrasting novel with master images, but it does not assume a subsequent memory effect in any form. Because the baseline GLM emerged as the optimal model from this first model space (see Section 4.1), it also formed the basis for all GLMs assuming a subsequent memory effect (see Table 1) by either adding a parametric modulator describing memory performance (*parametric models*; see Section 3.3) or separating the novelty regressor into different memory reports (*categorical models*; see Section 3.2).

### 3.2. Categorical memory models: two, three or five regressors

As the focus of our study was to optimize the fMRI modeling of the DM effect, we focused all our subsequent analyses on models, derived from the baseline GLM, that included at least one subsequent memory regressor. We first compared the following categorical GLMs:^2^

- Following the classic subsequent memory approach, stimuli can be grouped into two categories, later remembered and later forgotten, whereby *definitely old* and *probably old* responses are always considered remembered and *definitely new* and *probably new* responses are always categorized as forgotten. Neutral items with *unsure* responses can be either considered forgotten (*GLM_2-nf*) or remembered (*GLM_2-nr*) or randomly sampled as forgotten or remembered (*GLM_2-ns*), resulting in a model family with three models.
- Another option is to group novel images into three categories: *remembered* (responses 4-5), *forgotten* (responses 1-2), and *neutral* (response 3), yielding a model with three novelty regressors (*GLM_3*).
- When all five response types are considered, this leads to a model with five novelty regressors (*GLM_5*), which allows to model neural correlates of recognition, familiarity or recollection by applying the appropriate contrast vectors (see Düzel et al., 2011, Fig. 1A). A limitation of this model (as well as of the model using three regressors) was that not all subjects made use of all five response options during retrieval, such that this model could not be estimated for all subjects and results in ineffective data usage.

### 3.3. Parametric memory models: theoretical or empirical modulators

Instead of assuming categorical effects of memory performance, models can also account for a possible parametric effect, such that the observed activity follows the levels (or a function of the levels) of a parametric variable (here: memory rating). This is implemented by collecting all novel images into one onset regressor and adding a parametric modulator (PM) describing the assumed modulation of the trial-specific HRF by successful encoding as assessed with subsequent memory performance (see Table 1). In other words, these models add a trial-wise parametric regressor to the baseline GLM, which consists in a transformation of the subject’s subsequent memory responses. The transformation used by each model was either theoretically informed (see Figure 2A) or empirically inferred (see Figure 2B).

**Figure 2.**
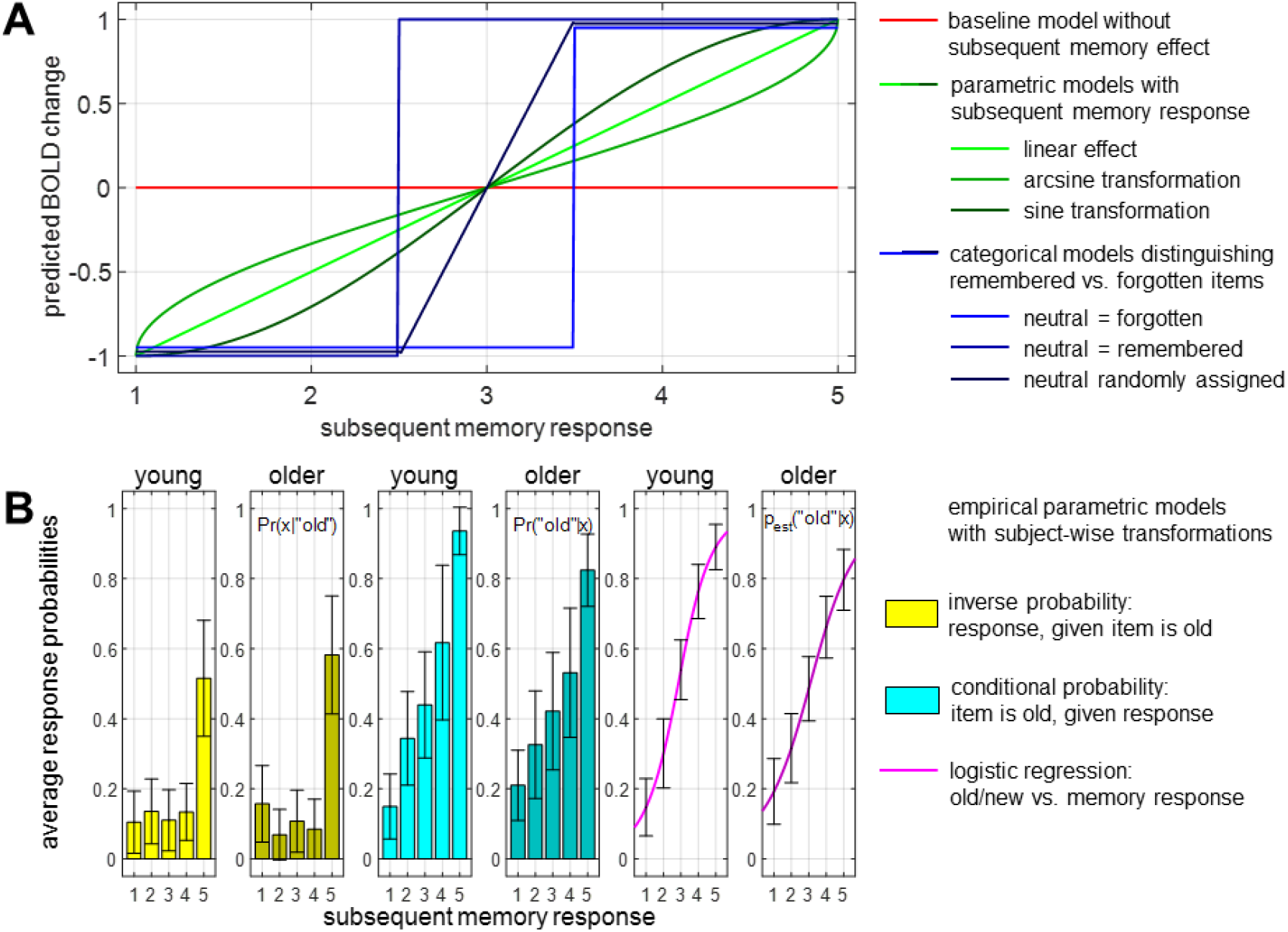
Parametric modulators for GLMs with subsequent memory effect. **(A)** Predicted signal change as a function of subsequent memory responses in the baseline GLM (red), the theoretical parametric GLMs (green) and the two-regressor categorical GLMs (blue). **(B)** Probabilities used as parametric modulators by empirical parametric GLMs. Error bars depict standard deviation (SD) across subjects; colors used in the plots correspond to box coloring in Supplementary Figure S2.

In the theoretical parametric models, a mathematical function of the subsequent memory report (*x*; responses 1-5) is applied to each item seen during the encoding session, yielding the parametric values modulating activity in the corresponding trials. Here, we implemented three plausible transformations:

- *GLM_1t-l*: a linear-parametric model; 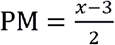 such that predicted activity increases linearly with memory response (see light green line in Figure 3A).
- *GLM_1t-a*: an arcsine-transformed parametric modulator; 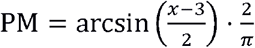; such that “sure” responses (definitely old/new) receive relatively higher weights than “unsure” responses (probably old/new) (see medium green line in Figure 3A).
- *GLM_1t-s*: a sine-transformed parametric modulator; 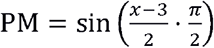; such that “unsure” responses receive relatively higher weights than “sure” responses (see dark green line in Figure 3A).

**Figure 3.**
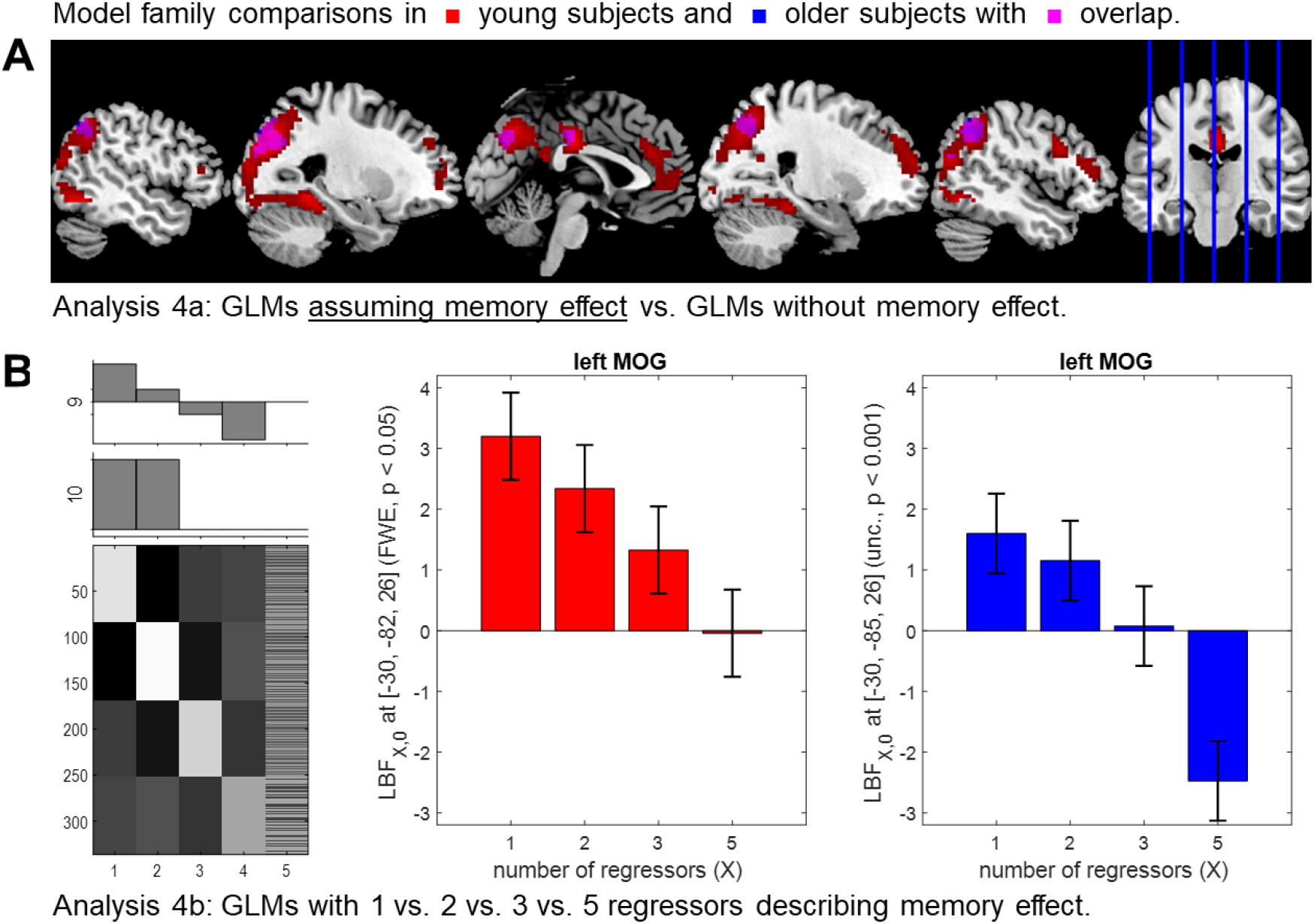
Effects of subsequent memory and number of regressors. **(A)** Selected-model maps in favor of GLMs modeling memory using one or two regressors, as obtained from young subjects (red), older subjects (blue) or both (magenta). Selected-model maps display model frequencies and color intensities range from 0 to 1. **(B)** Significant linear contrasts of the number of regressors used to describe memory (X) on the log Bayes (LBF) factor comparing models with X regressors against the baseline GLM, obtained in the global maxima of the respective conjunction contrasts, i.e. left middle occipital gyrus (MOG) in young subjects (red) and older subjects (blue). Bar plots depict contrasts of parameter estimates of the group-level model; error bars denote 90% confidence intervals (computed using SPM12).

All these transformations of *x* ∈ {1,2,3,4,5} ensure that – 1 ≤ PM ≤ +1, but they differ in their relative weighting of high confidence hits (5) and misses (1). In the linear-parametric model, the PM is proportional to *x*. The arcsine model puts a higher weight on definitely remembered (5) or forgotten (1) items compared with probably remembered (4) or forgotten (2) items, while the reverse is true for the sine model (see Figure 2A).

Alternatively, one can take a more data-driven approach and derive parametric modulators empirically from the behavioral data obtained in the retrieval session. To this end, all stimuli presented during retrieval, either *old* (i.e. previously seen during encoding) or *new*, are considered along with their corresponding memory reports (*x*; responses 1-5) to calculate probabilities which are then used as parametric modulators, e.g.:

- *GLM_1e-ip*: the inverse probability of subjects giving memory report *x*, given that an item was old, projected into the same range as above; PM = 2) · Pr(*x*|“old”) − 1;
- *GLM_1e-cp*: the conditional probability that at item was old, given memory report *x*, projected into the same range as above; PM = 2) · Pr “old”|*x*) − 1;
- *GLM_1e-lr*: in this model, logistic regression was used to predict whether a stimulus was old, given a subject’s memory report *x*, and the estimated posterior probability function was used as the parametric modulator, i.e. 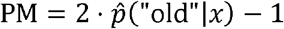

The resulting probabilities of all three models were normalized to the range – 1 PM +1 to ensure comparability with the theoretical parametric memory models. While the theoretical parametric GLMs are based on assumptions regarding the mapping of subsequent memory response to predicted BOLD signals (see Figure 2A), the empirical parametric GLMs incorporate subject-wise information, namely each subject’s response frequencies from the retrieval phase (see Figure 2B), which may improve model quality.

For all parametric GLMs, orthogonalization of parametric regressors was disabled in SPM, in order not to influence the estimates of the novelty onset regressor (Mumford et al., 2015).

## 4. Results

For each GLM, a cross-validated log model evidence (cvLME) map was calculated, and these maps were submitted to group-level cross-validated Bayesian model selection (cvBMS) analyses (see Section 2.5). Each analysis represents a specific modeling question, and each modeling question was separately addressed in young subjects (age ≤ 35, N = 92) and in older subjects (age ≥ 50, N = 153).

### 4.1. Effects of event duration, novelty and stimulus type

As a preliminary analysis step, we only considered the eight models without a subsequent memory effect, i.e. variations of the “baseline model” (see Section 3.1, Table 1). This allowed us to compare (i) point-event models vs. stimulus-duration models, to choose the optimal event duration, (ii) models that did or did not distinguish between novel and master images, to infer on the importance of the novelty effect in our models, and (iii) models that did or did not separate indoor and outdoor scenes, to assess the importance of considering this stimulus feature in an optimal model. Importantly, all of these analyses addressed model space dimensions of no interest. This means they served as sanity checks for logfile analysis and statistical modeling as well as validation of the memory paradigm (Düzel et al., 2011) and the cvBMS methodology (Soch et al., 2016).

First, we found that in both young and older participants, GLMs using an event duration of 2.5 s were preferred throughout the grey matter whereas white matter voxels are better described by GLMs using point events (see Supplementary Figure 3A). Presumably, this was an indirect result of the absence of task-related signal in white matter, such that simpler models (i.e., the GLMs assuming fewer processes) were selected automatically. Notably, the superiority of the trial duration models in grey matter was observed despite the fact that, due to the short inter-stimulus-intervals (see Section 2.2 and Figure 1C), regressors were more strongly correlated with each other when using a longer event duration.

Second, we observed that GLMs distinguishing between novel and master images outperformed GLMs not doing so throughout large portions of the occipital, parietal, and temporal lobes, extending into the bilateral parahippocampal cortex and hippocampus as well as the dorsolateral and rostral prefrontal cortex (see Supplementary Figure 3B), brain structures that are typically considered to constitute the human memory network (Jeong et al., 2015).

Third, cvBMS revealed that GLMs distinguishing between indoor and outdoor images outperformed GLMs not doing so in medial and lateral parts of the visual cortex (see Supplementary Figure 3C). Given the limited extent of clusters in the visual cortex favoring a separation of indoor and outdoor scenes and the aim of our study to optimize the modeling of the subsequent memory effect rather than perceptual processes, we decided not to include this additional modeling dimension.

Guided by the results of our preliminary analyses, we performed all memory-related model comparisons with GLMs using the actual trial length as event duration and separating images into novel and master, but not indoor and outdoor images.

### 4.2. Effects of subsequent memory and number of regressors

To address the effects of modeling subsequent memory on model quality, we calculated the log family evidence for all GLMs assuming any type of memory effect (categorical or parametric) and contrasted them against the log model evidence of the baseline GLM (assuming no memory effect). This analysis, i.e. identifying voxels in which models considering later memory collectively outperform the no-memory model, yielded considerably different results in young versus older subjects (see Figure 3): In young subjects, including a subsequent memory modulation led to an improved model fit in a set of brain regions that largely overlapped with those showing a superiority of the model family accounting for novelty (see Figure S3B and Section 4.1), including the dorsolateral prefrontal cortex (dlPFC), posterior cingulate cortex (PCC), precuneus (PreCun), lateral partietal cortices, portions of the ventral visual stream, and also the MTL (parahippocampal cortex and hippocampus). In older subjects, we observed qualitatively similar effects, but in a smaller number of voxels, and not in the dlPFC and parahippocampal cortex (see Figure 3A).

Among the GLMs modeling subsequent memory, we additionally tested for the influence of the number of regressors used to model the subsequent memory effect, which increases from the parametric memory models (1 parametric modulator per model; see Section 3.3) to the categorical memory models (2, 3 or 5 regressors; see Section 3.2). To this end, we calculated the LFE for each of these model families and subtracted the LME of the baseline GLM to compute log Bayes factors (LBF) maps in favor of memory models against a no-memory model. The rationale behind this was that some models assuming a subsequent memory effect might be too complex, essentially performing even worse than a model not accounting for memory performance at all. The LBF maps were then subjected to a one-way ANOVA model with the within-subject factor *number of regressors*, which has 4 levels (1, 2, 3, 5). There was a main effect of *number of regressors* throughout the whole brain (p < 0.05, FWE-corrected; results not shown). When performing a conjunction analysis between (i) a contrast of *GLMs_1* and *GLMs_2* against baseline and (ii) a t-contrast linearly decreasing with number of regressors, we found that the middle occipital gyrus (MOG), a brain structure with a previously demonstrated robust subsequent memory response (Kim, 2011), exhibited both a reliable DM effect as well as model quality gradients related to the number of regressors (see Figure 3B). These showed that only GLMs with one or two memory regressors outperformed the no-memory model whereas GLMs with three or five regressors were not significantly different from the null model or performed even worse, especially in the older subjects (see Figure 3B).

### 4.3. Parametric versus categorical subsequent memory models

The analyses described above indicate that parametric GLMs with one parametric modulator describing subsequent memory (*GLMs_1*) and categorical GLMs using two regressors for remembered vs. forgotten items (*GLMs_2*) perform best in regions previously implicated in successful memory formation (Kim, 2011). Treating these GLMs as model families, i.e. calculating log family evidences, and comparing the two families via group-level cvBMS, we observed a preference for parametric GLMs throughout the memory network (see Figure 4A), in regions largely overlapping with those that also showed a novelty effect (see Figure S3B and Section 4.1) and a memory effect (see Figure 3A and Section 4.2). The preference for parametric models could be observed in both age groups.

**Figure 4.**
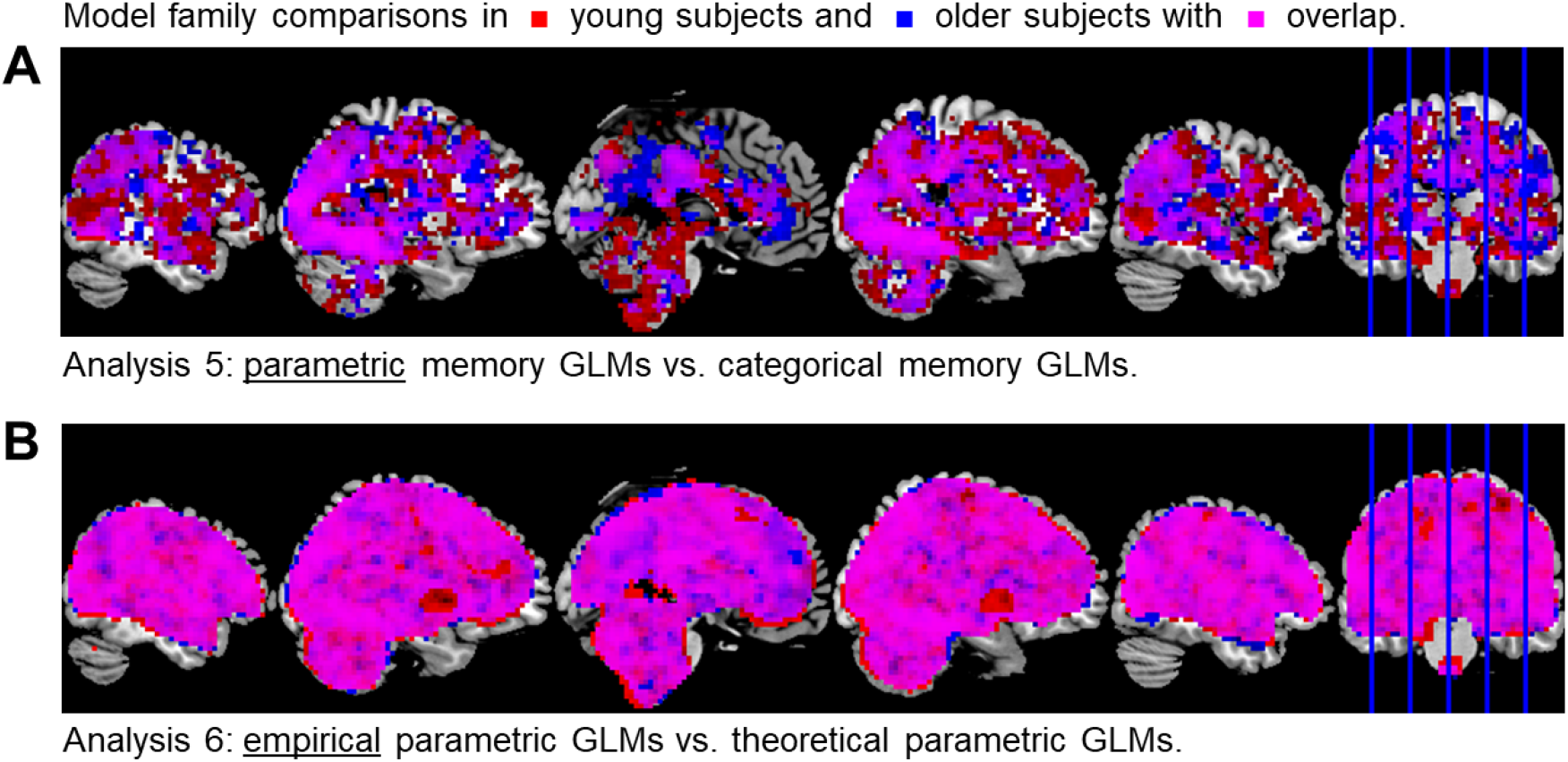
Parametric vs. categorical models of the subsequent memory effect. **(A)** Selected-model maps in favor of parametric GLMs against categorical GLMs. **(B)** Selected-model maps in favor of empirical parametric GLMs against theoretical parametric GLMs. Voxels displayed show the respective model preferences in young subjects (red) or older subjects (blue) or both groups (magenta).

Within the family of parametric memory models, we additionally compared theoretical GLMs (*GLMs_1t*) to empirical GLMs (*GLMs_1e*). Comparing these two sub-families via group-level cvBMS, we observed an almost whole-brain preference for the empirical GLMs (see Figure 4B).

### 4.4. Winning models within model families

The group-level results presented so far all refer to model families, i.e. sets of models whose collective quality was quantified via log family evidences calculated from log model evidences. This way, we have identified the three best performing families of GLMs: two-regressor categorical GLMs (*GLMs_2*), theoretical parametric GLMs (*GLMs_1t*), and empirical parametric GLMs (*GLMs_1e*). The final step of our model selection procedure was to test how models compared within these families, which was addressed by subjecting the respective cvLME maps to group-level cvBMS.

Within the *GLMs_2* family, the GLM categorizing neutral items with *don’t know* responses (3) as forgotten (*GLM_2-nf*) performed best in the majority of voxels (Figure 5A) when compared with the GLM categorizing those items as remembered (*GLM_2-nr*) or randomly distributing them among remembered and forgotten items (*GLM_2-ns*).

**Figure 5.**
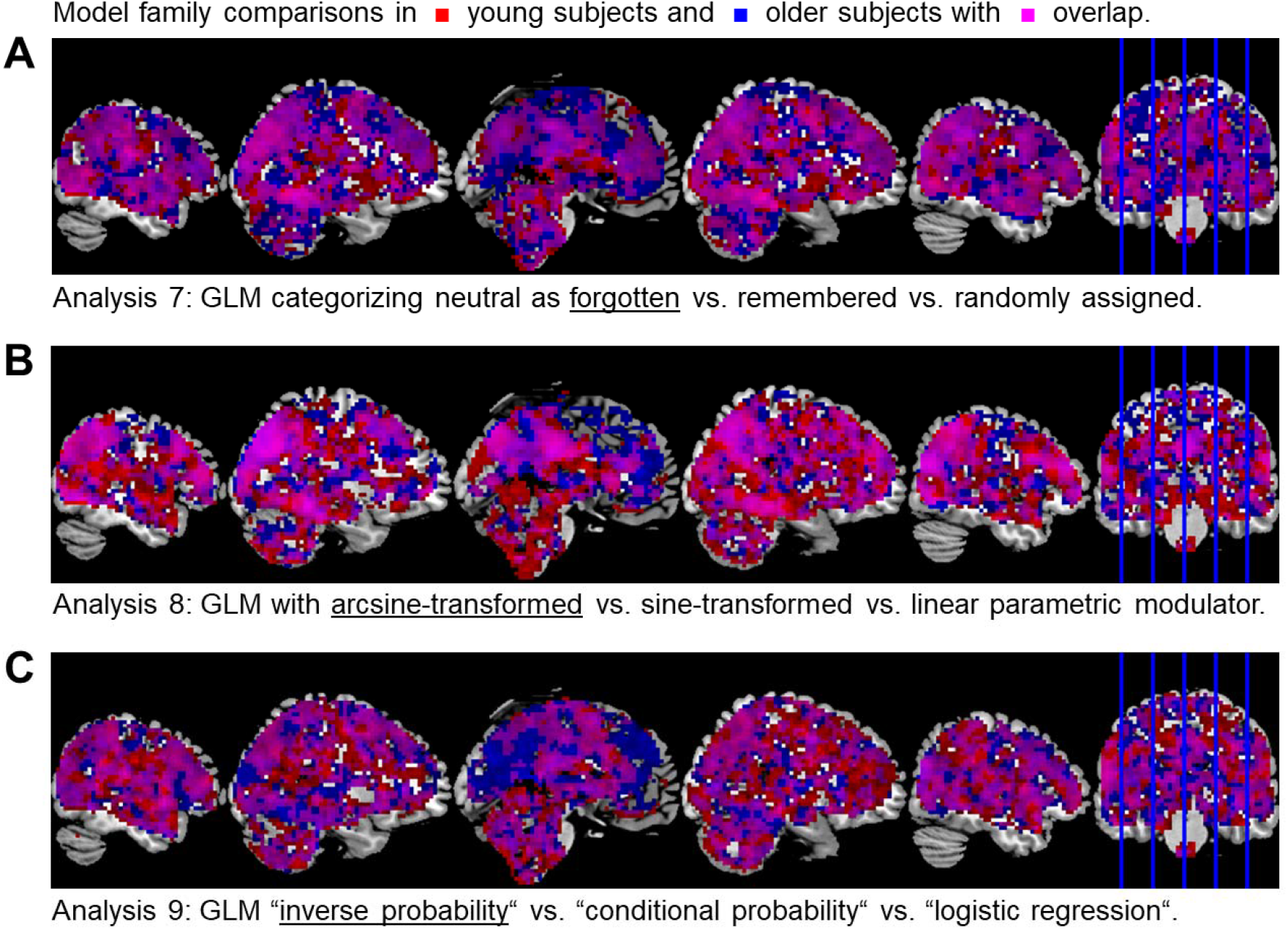
Winning models within model families. **(A)** Selected-model maps in favor of the GLM treating neutral images as forgotten items within the two-regressor categorical GLMs. **(B)** Selected-model maps favoring the GLM using an arcsine-transformed parametric modulator within the theoretical parametric GLMs. **(C)** Selected-model maps in favor of the GLM using an inverse probability parametric modulator within the empirical parametric GLMs. Voxels displayed show the respective model preferences in young subjects (red) or older subjects (blue) or both groups (magenta).

Within the *GLMs_1t* family, the GLM with the arcsine-transformed memory report as parametric modulator (*GLM_1t-a*) performed best in most voxels (Figure 5B) when compared with a sine (*GLM_1t-s*) or a linear (*GLM_1t-l*) transformation.

Within the *GLMs_1e* family, the GLM with the inverse probability Pr *x*|“old”) as parametric modulator (*GLM_1e-ip*) performed best in most voxels (Figure 5C) when compared with the conditional probability Pr “old”|*x*) (*GLM_1e-cp*) or logistic regression (*GLM_1e-lr*).

Overall, within-family differences were smaller than between-family differences, as indicated by lower likeliest frequencies (LFs) on the selected-model maps (cf. Figure 6 vs. Figure S3), reflecting more subtle modeling modifications within versus between families and age-related activation differences being larger in between-family comparisons.

**Figure 6.**
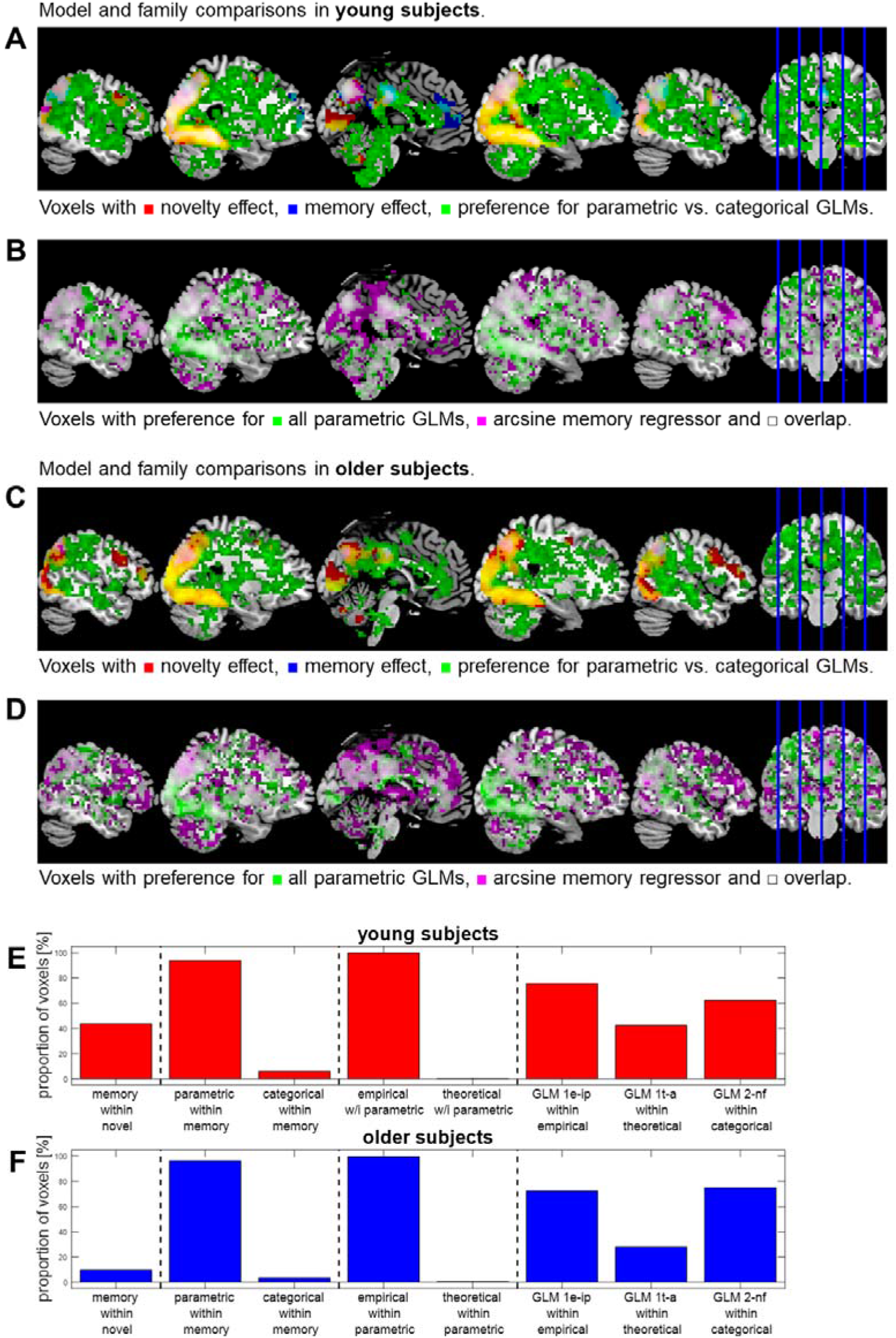
(see previous page). Model (family) comparisons (summary). **(A)** and **(B)** Selected-model maps (young subjects) in favor of GLMs assuming a novelty effect (red; see Figure S3B), a memory effect (blue; see Figure 3A), parametric vs. categorical memory effects (green; see Figure 4A) or an arcsine-shaped subsequent memory effect vs. other theoretical models (magenta; see Figure 5B). In most voxels with preference for parametric GLMs, there was also a preference for the arcsine model. **(C)** and **(D)** The corresponding selected-model maps from older subjects. **(E)** Proportion of voxels in which a model or family was selected (young subjects). “X within Y” is to be read as “probability that X was the selected family among voxels in which Y was the selected family”. **(F)** Same proportions as in E, obtained from older subjects.

### 4.5. Novelty and memory parameter estimates

The aforementioned analyses provide information about the models that best explain the BOLD signal in memory-related brain regions. They do, however, thus far not provide any information about the directionality, strength, or significance of the actual DM effect in the respective brain structures. To assess how the results of our model selection relate to group-level GLM results, we conducted second-level significance tests across the parameter estimates of the novelty and memory regressors from the three models identified as selected models in the three families that were performing best (see Figure 5). Replicating previous results (Kim, 2011; Maillet and Rajah, 2014), we observed memory-related activation differences in a temporo-parieto-occipital network and portions of the dlPFC (see Figure 7).^3^

**Figure 7.**
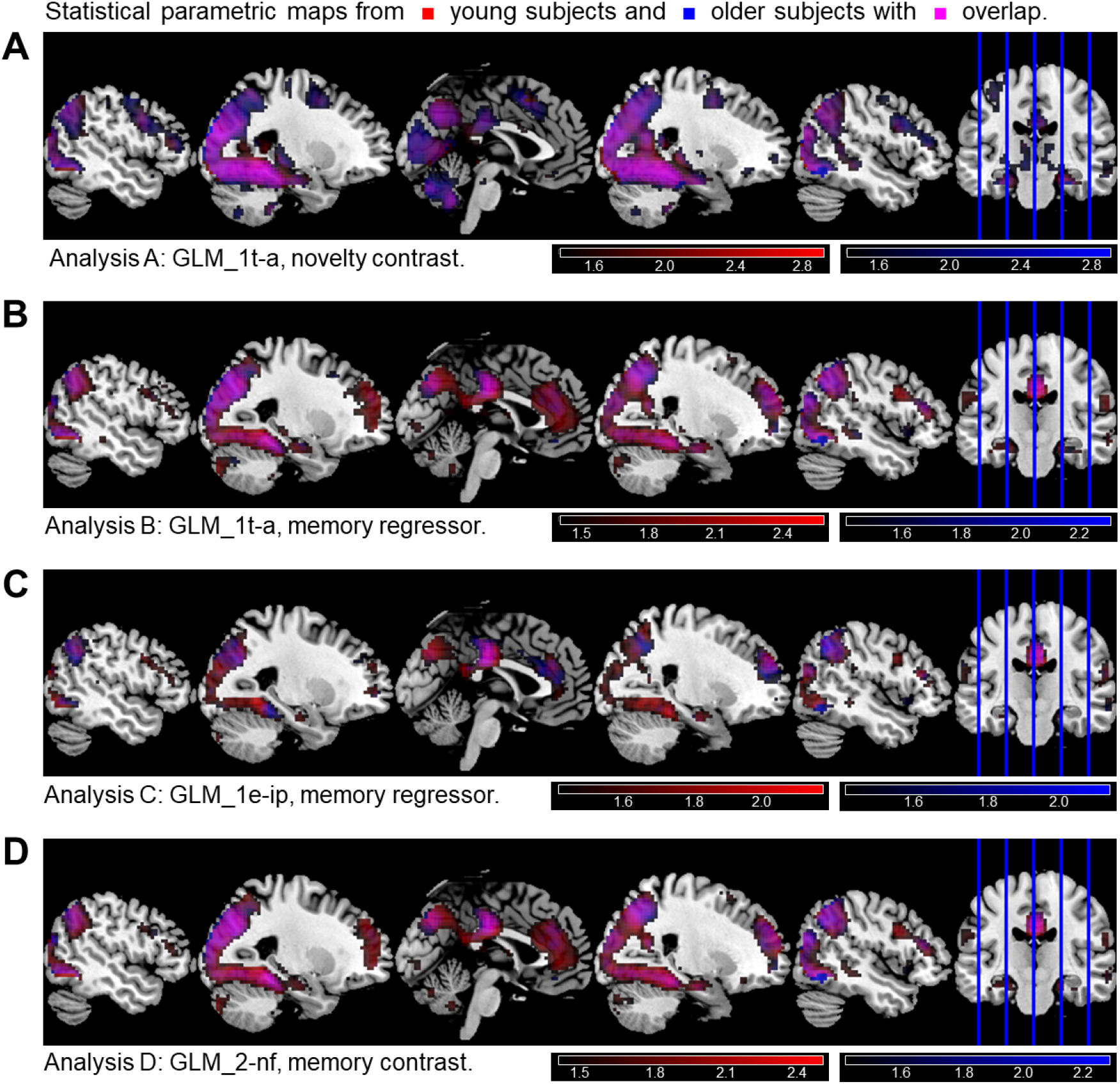
Exemplary statistical parametric maps. On the second level, a one-sample t-test was run across parameter estimates obtained from young subjects (red) and older subjects (blue) for **(A)** the novelty contrast (novelty vs. master images) and **(B)** the memory regressor of the theoretical-parametric GLM using the arcsine-transformed PM, **(C)** the memory regressor of the empirical-parametric GLM using the inverse probability PM and **(D)** the memory contrast (remembered vs. forgotten items) resulting from a two-regressor categorical GLM categorizing neutral responses as forgotten. In SPM, statistical inference was corrected for multiple comparisons (FWE, p < 0.05, k = 10), resulting in critical F-values for thresholding of SPMs (young: F > 27.01; older: F > 25.94). Color maps are scaled from the critical F-value to the maximum F-value in each map, in units of the decadic logarithm (see color bars).

In addition to this, we also report the subsequent memory performance as behavioral data (see Supplementary Table S2) and identify age-related differences with respect to response frequencies. Most prominently, older subjects significantly more often used high-confidence ratings and significantly less often used low-confidence ratings (see Table S2).

### 4.6. Replication in an independent cohort

Using the data from an independent replication cohort of young, healthy subjects (Assmann et al., 2020; Barman et al., 2014; Schott et al., 2014), we performed the analyses as described above. Performing these analyses using LME images from the additional cohort, we were largely able to replicate our results, sometimes with remarkable overlap between original and replication cohort (see Supplementary Figure S5), and sometimes with even stronger evidence for the most often selected model (see Supplementary Figure S8). Results from the replication cohort are displayed in Supplementary Figures S5-S10.

## 5. Discussion

We have applied cross-validated Bayesian model selection (Soch et al., 2016), a novel method for principled comparison between GLMs for fMRI data, to a previously described version of the subsequent memory paradigm (Düzel et al., 2011) in two large samples of young and older adults. By using the cvBMS approach, we have identified several ways to improve the modeling of subsequent memory effects in fMRI.

### 5.1. Optimal statistical modeling of subsequent memory effects

A key finding from our model selection was the preference of parametric over categorical GLMs of the fMRI subsequent memory effect (see Figures 3B, 4A and 6). At the model family level, GLMs with one memory regressor, a parametric modulator, outperformed GLMs with two, three or five memory regressors categorizing the events of interest. A core property of the cvLME approach is that it balances model accuracy and model complexity. With respect to our present analyses, this means that the categorical models allow for fitting more diverse activation patterns across memory reports, thereby achieving a higher accuracy when fitting the data. On the downside, their ability to generalize is rather limited, particularly when there is a low number of events in a given response category. In such cases, categorical models may fit tiny, but spurious irregularities between memory reports, indicating that they are not only more complex than necessary, but also prone to overfitting the data. On the other hand, parametric models are more parsimonious requiring only a single memory regressor, and thus are less likely to overfit the data.

An important caveat when using parametric models is the assumption of a parametric, or at least monotonic relationship between the parameter and the measured response, as often observed, for example when varying stimulus intensity or similar properties (Bogler et al., 2013; Soch et al., 2016, Fig. 3B; Soch et al., 2020, Fig. 8C). The question whether this assumption is met in the case of successful episodic memory encoding touches an intense debate in the memory research community that has been ongoing for decades. Several researchers have argued for a qualitative distinction of recollection and familiarity that is mirrored by a hierarchical architecture of the MTL memory system, with the hippocampus subserving context-rich, recollection-based memory, whereas rote, familiarity-based recognition memory relies on the perirhinal and parahippocampal cortices (Vargha-Khadem et al., 2001; Yonelinas et al., 2010). The alternative view emphasizes common processes in episodic and semantic memory and the high overlap between recollection and high-confidence familiarity, with activity of the MTL showing a quantitative rather than qualitative relationship with memory strength (Squire et al., 2007; Wixted and Squire, 2011).

The preference for parametric models observed in our cvBMS analysis seems, at first sight, to be more in line with the second view. It must, on the other hand, also be noted that, within the family of parametric memory models, non-linear transformations of subsequent memory performed better at describing the measured hemodynamic signals during memory encoding than a simple linear parametric modulation of the novelty regressor with memory confidence ratings. At the level of single models, the ones using the arcsine-transformed PM (theoretical) and the inverse probability PM (empirical) were favored by cvBMS. Both models put a high weight on stimuli recognized with high confidence (response “5”) relative to low-confidence recognition (response “4”). In the case of the inverse probability GLM, the group average (see Figure 2B) even suggests that the entire DM effect might by driven by a difference between high-confidence hits and all other conditions, which would essentially correspond to the recollection estimate proposed in the original publication of the paradigm used here (Düzel et al., 2011). In a supplementary analysis directly comparing the arcsine-transformed PM against the inverse probability PM, we found that model quality differences were rather unspecific within the human memory network, but that there were systematic age differences in cortical midline structures, with young subjects preferring the arcsine-transformed GLM and older subjects favoring the inverse probability GLM (see Supplementary Figure S4).

It must be emphasized that, even though the group average of the inverse probability PM is suggestive of a bias towards encoding predicting high-confidence memory, the very definition of this PM based on individual behavioral data allows for very different weighting of the response options at the level of single subjects. It can therefore also be employed in individuals with poor memory and largely absent recollection. On the downside, PMs with substantially different weighting might be difficult to compare at the group level. In this case, the model using the arcsine-transformed PM may be preferable.

### 5.2. Model preferences and age-related differences in the human memory network

Beyond the preference of a specific model of the DM effect, an overarching trend in our model selection results relates to the repeated observation of a distributed memory network in model preferences (see Figure 6A): When comparing regions with a novelty effect, regions with a subsequent memory effect and regions preferring parametric over categorical GLMs, there was a pronounced convergence of model preferences in multiple brain regions previously implicated in successful memory encoding (Kim, 2011), such as lateral and medial parietal cortices (e.g., PreCun), inferior temporal areas extending into the MTL with the hippocampus and parahippocampal cortex, as well as the dlPFC.

While there was overall convergence of brain regions exhibiting the same model preferences for novelty and subsequent memory across the entire study sample, this convergence was less pronounced in the older participants. On the one hand, there were almost no age differences in model preference regarding novelty (see Figure S3B), i.e. in both young and older participants, the model accounting for stimulus novelty significantly outperformed the model using a single regressor for all images. On the other hand, a considerable difference between age groups emerged with respect to subsequent memory effects (see Figure 3A). Here, the older participants showed a preference for models accounting for the DM effect in substantially fewer voxels and brain regions compared to the young group. Most likely because subsequent memory effects are generally weaker in older subjects, memory models also perform weaker when compared against the baseline GLM assuming no memory effect (see Figure 3B). In extreme cases, this can even mean that very complex models, such as the five-category GLM in the present study, may perform significantly worse than the memory null model, because the latter is prone to overfitting neural responses, which in turn decreases its generalizability.

We cannot exclude that even less complex categorical models may be inferior to a simple novelty-based model assuming no memory effect in participants with very poor memory, such as patients with early or pre-clinical Alzheimer’s disease. However, ongoing data analyses suggest that the DM effect may exhibit more extensive and robust differences between young and older individuals when compared to the novelty effect (J.S., A.R., J.K. and B.S., unpublished observations). In such situations, we suggest that the use of a relatively simple parametric model may provide a reasonable tradeoff between model complexity and utility.

### 5.3. Clinical implications and future directions

While episodic memory performance almost invariably declines during normal aging, accelerated memory decline is also a prominent symptom of Alzheimer’s disease (AD) (Buckner, 2004; Cansino, 2009; Rubin et al., 1998). Those observations at the behavioral level are mirrored by structural imaging findings showing age-related volume loss in the MTL (Raz et al., 2007) and pronounced MTL involvement in AD (Duara et al., 2008; Jack et al., 1998; Visser et al., 2002). To allow for early intervention, it is desirable to identify individuals developing AD at early clinical risk stages like subjective cognitive decline (SCD) or mild cognitive impairment (MCI) (Jessen et al., 2020). Considering the marked interindividual variability of age-related changes of encoding-related brain activity (Düzel et al., 2011), the DM paradigm might provide a useful tool in dissociating AD-related pathological changes from effects of normal aging. We suggest that the use of parametric models may help to further improve the utility of subsequent memory fMRI activations as a potential biomarker, as they are less dependent on individual memory performance when compared to categorical models.

### 5.4. Applicability beyond memory research

We have described the application of the cvBMS approach (Soch et al., 2016; Soch and Allefeld, 2018) to a paradigm that has previously been shown to elicit robust subsequent memory effects and is useful for detecting individual differences at the level of brain activity (Assmann et al., 2020; Barman et al., 2014; Düzel et al., 2011). While, to our knowledge, no previous study has employed our approach as extensively with respect to both model space and sample size, it should be emphasized that cvBMS should be applicable to essentially all cognitive paradigms in fMRI research that allow for multiple plausible first level models. As described above, we started our model selection procedure by assessing the influence of stimulus duration and content (i.e., indoor vs. outdoor), with both selections yielding clear preferences, which guided our subsequent analyses (see Figure S3). This provides further evidence for the utility of cvBMS in modeling decisions with respect to even very basic technical or stimulus-related aspects of a first-level fMRI model, similarly to its previously described application for deciding on the use of temporal and dispersion derivatives of the BOLD response (Soch et al., 2016). On the other hand, the cvBMS approach is not limited to such fundamental aspects of models, but can also be used to help decide between multiple models reflecting different theories of underlying cognitive and behavioral processes (Charpentier et al., 2020). The study by Charpentier and colleagues is particularly noteworthy from an Open Science perspective. The authors conducted a pre-registered study aimed at replication of their original findings. When their model of imitation learning from the first study could not be replicated, they employed cvBMS in an exploratory analysis and found a simpler model to be associated with higher exceedance probabilities in both their original data and the data of their pre-registered replication study. For future research, we suggest that pre-registered studies could also employ cvBMS and pre-register their complete model space, thereby allowing for more flexibility during data analysis, despite pre-registration.

In the present study, our interest was focused on the use of categorical versus parametric models, and on the model preferences within the respective model families. Parametric regressors with more than two values are commonly coded as linear scales (Heinzel et al., 2005; Northoff et al., 2009) and, in the present study, a linear scale of recognition confidence was used in our default parametric model. We additionally employed an arcsine-transformed scale, which was inspired by the use of arcsine transformations for proportion data in statistics (Hernández et al., 2018; Lin and Xu, 2020). The choice of arcsine over other inverse sigmoid transformations like the logit or Fisher z-transformations was due to the non-asymptotic nature of the arcsine function. We are aware that other transformations with similar shapes (e.g., a cubic function) will likely yield similar results. While such differences are probably negligible due to the rather coarse five-step scaling of our parametric modulators, the shape of the transformation might become more important when employing scales with a higher resolution, such as visual analog scales (Northoff et al., 2009). In such cases, comparing parametric models with several alternative transformations may be helpful to further improve the model fit.

### 5.5. Limitations

One limitation of the present approach is that any parametric model assumes an at least monotonic relationship between memory confidence and brain activation patterns. Evidently, such a relationship is plausible for any model assuming increasing memory strength as a function of increasing MTL engagement (Wixted and Squire, 2011), but it can also be applicable to hierarchical models of memory performance when considering, for example, that recollection is highly correlated with high memory confidence and accompanied by an additional familiarity signal (Yonelinas et al., 2010). However, caution is necessary as confidence and recognition accuracy may not necessarily be correlated under all circumstances (Busey et al., 2000). Furthermore, the assumption of a monotonic relationship will likely be violated when applying single-process models that also include implicit memory processes like priming (Berry et al., 2012). For example, previous studies have demonstrated encoding-related activations predicting explicit memory, but de-activations predicting priming in the fusiform gyrus (Schott et al., 2006), and a possibly reverse pattern in the right temporo-parietal junction (Schott et al., 2006; Uncapher and Wagner, 2009; Wimber et al., 2010).

### 5.6. Conclusions

Our results suggest that a systematic model selection approach favors parametric over categorical models in first-level GLM-based analysis of the fMRI subsequent memory effect.

While it would be, in our view, premature to draw a conclusion with respect to hierarchical versus single-process models of explicit memory function in the human memory network based on these results, our results do provide a strong rationale for the use of parametric models in studies focusing on between-group differences, particularly in older humans and individuals with impaired memory performance.

## Supporting information

Supplementary Methods, Results, and Figures

## 6. Notes

## 6.1. Acknowledgments

The authors would like to thank Kerstin Möhring, Katja Neumann, Ilona Wiedenhöft, and Claus Tempelmann for assistance with MRI acquisition.

## 6.2. Author contributions

Conceptualization: J.S., A.R., J.M.K., A.R.-K., E.D., B.H.S.; Data curation: J.S., A.R., L.K., M.R., A.S., B.H.S.; Formal analysis: J.S., A.R., J.M.K., B.H.S.; Funding acquisition: A.R., A.R.-K., E.D., B.H.S.; Investigation: A.R., A.A., L.K., M.R., A.S.; Methodology: J.S.; Project administration: A.R., E.D., B.H.S.; Resources: E.D.; Software: J.S., H.S.; Supervision: A.R., E.D., B.H.S.; Validation: J.S., A.R., A.A., B.H.S.; Visualization: J.S., A.R.; Writing-original draft: J.S., A.R., J.M.K., B.H.S.; Writing - review & editing: J.S., A.R., J.M.K., A.M., G.Z., E.D., B.H.S.

## 6.3. Data Availability Statement

Due to data protection concerns, sharing of the entire data set underlying this study is not possible at the moment. However, we provide cross-validated log model evidence (cvLME) maps for 19 models (see Table 1) as well as general linear model (GLM) contrast images for 4 parameters (see Figure 7) from all 245 subjects underlying the group analyses reported in this paper as NeuroVault collections (https://neurovault.org/collections/BYSHJNCO/, https://neurovault.org/collections/FVUWBMVP). MATLAB code and instructions to process these data can be found in an accompanying GitHub repository (https://github.com/JoramSoch/FADE_BMS).

## Footnotes

1 In future studies, the data from the young participants of the present study will serve as baseline data to investigate effects of aging and neurodegeneration.

2 Note that, from here on, the first number after “GLM” in a model name corresponds to the number of regressors used to describe the subsequent memory effect (see Figure 2B), i.e. *GLM_0\** = no memory regressor; *GLM_1\** = one memory regressor; *GLM_2\** = two memory regressors; etc.

3 Please note that the analyses of the DM effect were limited to F-contrasts in order to verify the overall applicability of the winning models. Detailed analyses of the subsequent memory effects, with a particular focus on age-related differences, are beyond the scope of the current study and will be reported elsewhere.

## References

Assmann A, Richter A, Schütze H, Soch J, Barman A, Behnisch G, Knopf L, Raschick M, Schult A, Wüstenberg T, Behr J, Düzel E, Seidenbecher CI, Schott BH. 2020. Neurocan genome[wide psychiatric risk variant affects explicit memory performance and hippocampal function in healthy humans. Eur J Neurosci ejn.14872. doi:10.1111/ejn.14872

Bainbridge WA, Berron D, Schütze H, Cardenas-Blanco A, Metzger C, Dobisch L, Bittner D, Glanz W, Spottke A, Rudolph J, Brosseron F, Buerger K, Janowitz D, Fliessbach K, Heneka M, Laske C, Buchmann M, Peters O, Diesing D, Li S, Priller J, Spruth EJ, Altenstein S, Schneider A, Kofler B, Teipel S, Kilimann I, Wiltfang J, Bartels C, Wolfsgruber S, Wagner M, Jessen F, Baker CI, Düzel E. 2019. Memorability of photographs in subjective cognitive decline and mild cognitive impairment: Implications for cognitive assessment. Alzheimer’s Dement Diagnosis, Assess Dis Monit 11:610–618. doi:10.1016/j.dadm.2019.07.005

Barman A, Assmann A, Richter S, Soch J, SchÃ¼tze H, WÃ¼stenberg T, Deibele A, Klein M, Richter A, Behnisch G, DÃ¼zel E, Zenker M, Seidenbecher CI, Schott BH. 2014. Genetic variation of the RASGRF1 regulatory region affects human hippocampus-dependent memory. Front Hum Neurosci 8:1–12. doi:10.3389/fnhum.2014.00260

Berry CJ, Shanks DR, Speekenbrink M, Henson RNA. 2012. Models of recognition, repetition priming, and fluency: Exploring a new framework. Psychol Rev 119:40–79. doi:10.1037/a0025464

Betts MJ, Cardenas-Blanco A, Kanowski M, Spottke A, Teipel SJ, Kilimann I, Jessen F, Düzel E. 2019. Locus coeruleus MRI contrast is reduced in Alzheimer’s disease dementia and correlates with CSF Aβ levels. Alzheimer’s Dement Diagnosis, Assess Dis Monit 11:281–285. doi:10.1016/j.dadm.2019.02.001

Bodnar M, Achim AM, Malla AK, Joober R, Benoit A, Lepage M. 2012. Functional magnetic resonance imaging correlates of memory encoding in relation to achieving remission in first-episode schizophrenia. Br J Psychiatry 200:300–307. doi:10.1192/bjp.bp.111.098046

Bogler C, Bode S, Haynes J-D. 2013. Orientation pop-out processing in human visual cortex. Neuroimage 81:73–80. doi:10.1016/j.neuroimage.2013.05.040

Brewer JB. 1998. Making Memories: Brain Activity that Predicts How Well Visual Experience Will Be Remembered. Science (80-) 281:1185–1187. doi:10.1126/science.281.5380.1185

Buckner RL. 2004. Memory and Executive Function in Aging and AD. Neuron 44:195–208. doi:10.1016/j.neuron.2004.09.006

Busey TA, Tunnicliff J, Loftus GR, Loftus EF. 2000. Accounts of the confidence-accuracy relation in recognition memory. Psychon Bull Rev 7:26–48. doi:10.3758/BF03210724

Cansino S. 2009. Episodic memory decay along the adult lifespan: A review of behavioral and neurophysiological evidence. Int J Psychophysiol 71:64–69. doi:10.1016/j.ijpsycho.2008.07.005

Charpentier CJ, Iigaya K, O’Doherty JP. 2020. A Neuro-computational Account of Arbitration between Choice Imitation and Goal Emulation during Human Observational Learning. Neuron 106:687–699.e7. doi:10.1016/j.neuron.2020.02.028

Daselaar SM, Fleck MS, Dobbins IG, Madden DJ, Cabeza R. 2006. Effects of healthy aging on hippocampal and rhinal memory functions: An event-related fMRI study. Cereb Cortex 16:1771–1782. doi:10.1093/cercor/bhj112

Davachi L, Mitchell JP, Wagner AD. 2003. Multiple routes to memory: Distinct medial temporal lobe processes build item and source memories. Proc Natl Acad Sci 100:2157–2162. doi:10.1073/pnas.0337195100

Dennis NA, Hayes SM, Prince SE, Madden DJ, Huettel SA, Cabeza R. 2008. Effects of aging on the neural correlates of successful item and source memory encoding. J Exp Psychol Learn Mem Cogn 34:791–808. doi:10.1037/0278-7393.34.4.791

Duara R, Loewenstein DA, Potter E, Appel J, Greig MT, Urs R, Shen Q, Raj A, Small B, Barker W, Schofield E, Wu Y, Potter H. 2008. Medial temporal lobe atrophy on MRI scans and the diagnosis of Alzheimer disease. Neurology 71:1986–1992. doi:10.1212/01.wnl.0000336925.79704.9f

Düzel E, Acosta[Cabronero J, Berron D, Biessels GJ, Björkman[Burtscher I, Bottlaender M, Bowtell R, Buchem M v., Cardenas[Blanco A, Boumezbeur F, Chan D, Clare S, Costagli M, Rochefort L, Fillmer A, Gowland P, Hansson O, Hendrikse J, Kraff O, Ladd ME, Ronen I, Petersen E, Rowe JB, Siebner H, Stoecker T, Straub S, Tosetti M, Uludag K, Vignaud A, Zwanenburg J, Speck O. 2019. European Ultrahigh[Field Imaging Network for Neurodegenerative Diseases (EUFIND). Alzheimer’s Dement Diagnosis, Assess Dis Monit 11:538–549. doi:10.1016/j.dadm.2019.04.010

Düzel E, Schütze H, Yonelinas AP, Heinze H-J. 2011. Functional phenotyping of successful aging in long-term memory: Preserved performance in the absence of neural compensation. Hippocampus 21:803–814. doi:10.1002/hipo.20834

Fernández G, Weyerts H, Schrader-Bölsche M, Tendolkar I, Smid HGOM, Tempelmann C, Hinrichs H, Scheich H, Elger CE, Mangun GR, Heinze HJ. 1998. Successful verbal encoding into episodic memory engages the posterior hippocampus: A parametrically analyzed functional magnetic resonance imaging study. J Neurosci 18:1841–1847. doi:10.1523/jneurosci.18-05-01841.1998

Fletcher P, Stephenson C, Carpenter T, Donovan T, Bullmore E. 2003. Regional Brain Activations Predicting Subsequent Memory Success: An Event-Related Fmri Study of the Influence of Encoding Tasks. Cortex 39:1009–1026. doi:10.1016/S0010-9452(08)70875-X

Friston KJ, Holmes AP, Worsley KJ, Poline J-P, Frith CD, Frackowiak RSJ. 1994. Statistical parametric maps in functional imaging: A general linear approach. Hum Brain Mapp 2:189–210. doi:10.1002/hbm.460020402

Heinzel A, Bermpohl F, Niese R, Pfennig A, Pascual-Leone A, Schlaug G, Northoff G. 2005. How do we modulate our emotions? Parametric fMRI reveals cortical midline structures as regions specifically involved in the processing of emotional valences. Cogn Brain Res 25:348–358. doi:10.1016/j.cogbrainres.2005.06.009

Henson RN, Rugg MD, Shallice T, Josephs O, Dolan RJ. 1999. Recollection and familiarity in recognition memory: an event-related functional magnetic resonance imaging study. J Neurosci 19:3962–72.

Hernández MÁ, Andrés AM, Tejedor IH. 2018. One-tailed asymptotic inferences for the difference of proportions: Analysis of 97 methods of inference. J Biopharm Stat 28:1090–1104. doi:10.1080/10543406.2018.1452028

Hinrichs H, Scholz M, Tempelmann C, Woldorff MG, Dale AM, Heinze H-J. 2000. Deconvolution of event-related fMRI responses in fast-rate experimental designs: tracking amplitude variations. J Cogn Neurosci 12:76–89. doi:10.1162/089892900564082

Jack CR, Petersen RC, Xu Y, O’Brien PC, Smith GE, Ivnik RJ, Tangalos EG, Kokmen E. 1998. Rate of medial temporal lobe atrophy in typical aging and Alzheimer’s disease. Neurology 51:993–999. doi:10.1212/WNL.51.4.993

Jeong W, Chung CK, Kim JS. 2015. Episodic memory in aspects of large-scale brain networks. Front Hum Neurosci 9:1–15. doi:10.3389/fnhum.2015.00454

Jessen F, Amariglio RE, Buckley RF, van der Flier WM, Han Y, Molinuevo JL, Rabin L, Rentz DM, Rodriguez-Gomez O, Saykin AJ, Sikkes SAM, Smart CM, Wolfsgruber S, Wagner M. 2020. The characterisation of subjective cognitive decline. Lancet Neurol 19:271–278. doi:10.1016/S1474-4422(19)30368-0

Jessen F, Spottke A, Boecker H, Brosseron F, Buerger K, Catak C, Fliessbach K, Franke C, Fuentes M, Heneka MT, Janowitz D, Kilimann I, Laske C, Menne F, Nestor P, Peters O, Priller J, Pross V, Ramirez A, Schneider A, Speck O, Spruth EJ, Teipel S, Vukovich R, Westerteicher C, Wiltfang J, Wolfsgruber S, Wagner M, Düzel E. 2018. Design and first baseline data of the DZNE multicenter observational study on predementia Alzheimer’s disease (DELCODE). Alzheimer’s Res Ther 10:1–10. doi:10.1186/s13195-017-0314-2

Kim H. 2011. Neural activity that predicts subsequent memory and forgetting: A meta-analysis of 74 fMRI studies. Neuroimage 54:2446–2461. doi:10.1016/j.neuroimage.2010.09.045

Kim H, Cabeza R. 2007. Trusting our memories: dissociating the neural correlates of confidence in veridical versus illusory memories. J Neurosci 27:12190–7. doi:10.1523/JNEUROSCI.3408-07.2007

Likert R. 1932. A technique for the measurement of attitudes. Arch Psychol 22:55.

Lin L, Xu C. 2020. Arcsine[based transformations for meta[analysis of proportions: Pros, cons, and alternatives. Heal Sci Reports 3:1–6. doi:10.1002/hsr2.178

Maillet D, Rajah MN. 2014. Age-related differences in brain activity in the subsequent memory paradigm: A meta-analysis. Neurosci Biobehav Rev 45:246–257. doi:10.1016/j.neubiorev.2014.06.006

Mumford JA, Poline J-B, Poldrack RA. 2015. Orthogonalization of Regressors in fMRI Models. PLoS One 10:e0126255. doi:10.1371/journal.pone.0126255

Northoff G, Schneider F, Rotte M, Matthiae C, Tempelmann C, Wiebking C, Bermpohl F, Heinzel A, Danos P, Heinze HJ, Bogerts B, Walter M, Panksepp J. 2009. Differential parametric modulation of self-relatedness and emotions in different brain regions. Hum Brain Mapp 30:369–382. doi:10.1002/hbm.20510

Otten LJ, Rugg MD. 2001. Electrophysiological correlates of memory encoding are task-dependent. Cogn Brain Res 12:11–18. doi:10.1016/S0926-6410(01)00015-5

Paller KA, Kutas M, Mayes AR. 1987. Neural correlates of encoding in an incidental learning paradigm. Electroencephalogr Clin Neurophysiol 67:360–371. doi:10.1016/0013-4694(87)90124-6

Raz N, Rodrigue KM, Haacke EM. 2007. Brain Aging and Its Modifiers: Insights from in Vivo Neuromorphometry and Susceptibility Weighted Imaging. Ann N Y Acad Sci 1097:84–93. doi:10.1196/annals.1379.018

Richardson MP, Strange BA, Duncan JS, Dolan RJ. 2003. Preserved verbal memory function in left medial temporal pathology involves reorganisation of function to right medial temporal lobe. Neuroimage 20:S112–S119. doi:10.1016/j.neuroimage.2003.09.008

Richter A, Barman A, Wüstenberg T, Soch J, Schanze D, Deibele A, Behnisch G, Assmann A, Klein M, Zenker M, Seidenbecher C, Schott BH. 2017. Behavioral and neural manifestations of reward memory in carriers of low-expressing versus high-expressing genetic variants of the dopamine D2 receptor. Front Psychol 8:1–13. doi:10.3389/fpsyg.2017.00654

Rubin EH, Storandt M, Miller JP, Kinscherf DA, Grant EA, Morris JC, Berg L. 1998. A prospective study of cognitive function and onset of Dementia in cognitively healthy elders. Arch Neurol 55:395–401. doi:10.1001/archneur.55.3.395

Schoemaker D, Gauthier S, Pruessner JC. 2014. Recollection and Familiarity in Aging Individuals with Mild Cognitive Impairment and Alzheimer’s Disease: A Literature Review. Neuropsychol Rev 24:313–331. doi:10.1007/s11065-014-9265-6

Schott BH, Assmann A, Schmierer P, Soch J, Erk S, Garbusow M, Mohnke S, Pöhland L, Romanczuk-Seiferth N, Barman A, Wüstenberg T, Haddad L, Grimm O, Witt S, Richter S, Klein M, Schütze H, Mühleisen TW, Cichon S, Rietschel M, Noethen MM, Tost H, Gundelfinger ED, Düzel E, Heinz A, Meyer-Lindenberg A, Seidenbecher CI, Walter H. 2014. Epistatic interaction of genetic depression risk variants in the human subgenual cingulate cortex during memory encoding. Transl Psychiatry 4:e372–e372. doi:10.1038/tp.2014.10

Schott BH, Richardson-Klavehn A, Henson RNA, Becker C, Heinze H-J, Düzel E. 2006. Neuroanatomical dissociation of encoding processes related to priming and explicit memory. J Neurosci 26:792–800. doi:10.1523/JNEUROSCI.2402-05.2006

Soch J, Allefeld C. 2018. MACS – a new SPM toolbox for model assessment, comparison and selection. J Neurosci Methods 306:19–31. doi:10.1016/j.jneumeth.2018.05.017

Soch J, Allefeld C, Haynes JD. 2020. Inverse transformed encoding models – a solution to the problem of correlated trial-by-trial parameter estimates in fMRI decoding. Neuroimage 209:116449. doi:10.1016/j.neuroimage.2019.116449

Soch J, Haynes J-D, Allefeld C. 2016. How to avoid mismodelling in GLM-based fMRI data analysis: cross-validated Bayesian model selection. Neuroimage 141:469–489. doi:10.1016/j.neuroimage.2016.07.047

Squire LR, Wixted JT, Clark RE. 2007. Recognition memory and the medial temporal lobe: a new perspective. Nat Rev Neurosci 8:872–883. doi:10.1038/nrn2154

Teipel SJ, Metzger CD, Brosseron F, Buerger K, Brueggen K, Catak C, Diesing D, Dobisch L, Fliebach K, Franke C, Heneka MT, Kilimann I, Kofler B, Menne F, Peters O, Polcher A, Priller J, Schneider A, Spottke A, Spruth EJ, Thelen M, Thyrian RJ, Wagner M, Düzel E, Jessen F, Dyrba M. 2018. Multicenter Resting State Functional Connectivity in Prodromal and Dementia Stages of Alzheimer’s Disease. J Alzheimer’s Dis 64:801–813. doi:10.3233/JAD-180106

Towgood K, Barker GJ, Caceres A, Crum WR, Elwes RDC, Costafreda SG, Mehta MA, Morris RG, von Oertzen TJ, Richardson MP. 2015. Bringing memory fMRI to the clinic: Comparison of seven memory fMRI protocols in temporal lobe epilepsy. Hum Brain Mapp 36:1595–1608. doi:10.1002/hbm.22726

Turk-Browne NB, Yi DJ, Chun MM. 2006. Linking implicit and explicit memory: Common encoding factors and shared representations. Neuron 49:917–927. doi:10.1016/j.neuron.2006.01.030

Uncapher MR, Wagner AD. 2009. Posterior parietal cortex and episodic encoding: Insights from fMRI subsequent memory effects and dual-attention theory. Neurobiol Learn Mem 91:139–154. doi:10.1016/j.nlm.2008.10.011

Vargha-Khadem F, Gadian DG, Mishkin M. 2001. Dissociations in cognitive memory: the syndrome of developmental amnesia. Philos Trans R Soc London Ser B Biol Sci 356:1435–1440. doi:10.1098/rstb.2001.0951

Visser PJ, Verhey FRJ, Hofman PAM, Scheltens P, Jolles J. 2002. Medial temporal lobe atrophy predicts Alzheimer’s disease in patients with minor cognitive impairment. J Neurol Neurosurg Psychiatry 72:491–497. doi:10.1136/jnnp.72.4.491

Wagner AD, Schacter DL, Rotte M, Koutstaal W, Maril A, Dale AM, Rosen BR, Buckner RL. 1998. Building memories: Remembering and forgetting of verbal experiences as predicted by brain activity. Science (80-) 281:1188–1191. doi:10.1126/science.281.5380.1188

Wimber M, Heinze H-J, Richardson-Klavehn A. 2010. Distinct frontoparietal networks set the stage for later perceptual identification priming and episodic recognition memory. J Neurosci 30:13272–13280. doi:10.1523/JNEUROSCI.0588-10.2010

Wixted JT, Squire LR. 2011. The medial temporal lobe and the attributes of memory. Trends Cogn Sci 15:210–217. doi:10.1016/j.tics.2011.03.005

World Medical Association. 2013. World Medical Association Declaration of Helsinki: Ethical principles for medical research involving human subjects. JAMA 310:2191–2194. doi:10.1001/jama.2013.281053

Yonelinas AP. 1994. Receiver-operating characteristics in recognition memory: Evidence for a dual-process model. J Exp Psychol Learn Mem Cogn 20:1341–1354. doi:10.1037/0278-7393.20.6.1341

Yonelinas AP, Aly M, Wang W-C, Koen JD. 2010. Recollection and familiarity: Examining controversial assumptions and new directions. Hippocampus 20:1178–1194. doi:10.1002/hipo.20864

Zierhut K, Bogerts B, Schott B, Fenker D, Walter M, Albrecht D, Steiner J, Schütze H, Northoff G, Düzel E, Schiltz K. 2010. The role of hippocampus dysfunction in deficient memory encoding and positive symptoms in schizophrenia. Psychiatry Res Neuroimaging 183:187–194. doi:10.1016/j.pscychresns.2010.03.007

