## Supplementary Methods, Results, and Figures for "Bayesian model selection favors parametric over categorical fMRI subsequent memory models in young and older adults"

##### Power calculation and sample size

The data were acquired as part of a study aimed at the comprehensive identification and characterization of biomarkers that predict successful versus accelerated aging of the human memory systems. Sample sizes were computed using G*Power (Faul et al., 2007). Sample size estimation was aimed at detecting moderately strong effects (Cohen's d = 0.6) in between-subjects t-contrasts with a significance level at p < .001, the recommended *a priori* significance level for cluster-based family-wise error (FWE) correction (Eklund et al., 2016). With an allocation ratio of *older* : *young* = 3 : 2, the power analysis yielded a minimum sample of 82 young and 122 older subjects. Data collection is ongoing to ultimately yield twice the calculated sample size, thus allowing for further subgroup-based analyses in the future.

##### Comparison of winning models across model families

In the main analysis, we identified winning models in the three best-fitting GLM families (see Section 4.4 and Figure 5). In a supplementary analysis, we compared the winning models from the two parametric families, the empirical parametric GLM using the inverse probability $\mathrm{Pr}(x|\text{"old"})$ as parametric modulator (PM) (see Figure 5C) and the theoretical parametric GLM with the arcsine-transformed PM (see Figure 5B). To this end, the LME images associated with these two models were subjected to group-level BMS (see Section 2.5) and selected-model maps were generated.

##### Replication of results from young subjects in independent cohort

The fMRI data analyzed in the present study came from a cohort consisting of 106 young and 153 older subjects who performed a subsequent memory paradigm previously termed the FADE task (functional activity deviation during encoding; Düzel et al., 2011). The same task, albeit with slightly different trial timings, has previously been employed in an earlier study (Schott et al., 2014; Assmann et al., 2020; henceforth termed "yFADE"), yielding another data set of 117 young subjects (60 male, 57 female, age range 19-33, mean age 24.37 ± 2.595 years). Here, we used this data set for an independent replication of the results obtained in the young subjects of the present study. As there was no overlap among study participants between the group, the additional cohort can be considered a biological replicate (Blainey et al., 2014).

The data from this replication cohort differed only marginally in terms of fMRI acquisition parameters (for details, see Assmann et al., 2020) and stimulus timing settings (see Figure S1).

To maximize comparability, the replication data, which had been acquired at a higher spatial resolution, were normalized to the same voxel size and smoothed with the same kernel width as the original data (see Table S1).

### Supplementary Results

#### Behavioral results

Relative response frequencies for all five confidence levels as well as hit rate (the proportion of old items rated as 4 or 5), false alarm rate (the proportion of new items rated as 4 or 5) and corrected hit rate (hits minus false alarms) are summarized in Table S2, separated by cohort (young subjects, older subjects, replication cohorts). Two-sample t-tests comparing the response frequencies of young and older subjects for each confidence level indicated that, compared to young subjects, older subjects often used high-confidence ratings (1 and 5) significantly more and significantly less often used low-confidence ratings (2 and 4) for stimuli presented during encoding. Furthermore, older subjects displayed a significantly higher false alarm rate and a significantly lower corrected hit rate.

#### Comparison of winning models across model families

When comparing the two best-fitting parametric GLMs against each other, i.e. when comparing *GLM_1t-a* vs. *GLM_1e-ip*, we observed rather unsystematic model preferences in the human memory network (see Figure S4, 2nd and 4th section from left), but a relatively systematic age difference in memory-related midline structures (see Figure S4, 3rd section from left): While young subjects represent subsequent memory ratings according to the theoretical GLM with the arcsine-transformed PM (see Figure S4B) in the precuneus (PreCun), posterior cingulate cortex (PCC) and medial prefrontal cortex (mPFC), older subjects show a model preference for the empirical GLM with the inverse probability PM (see Figure S4A).

#### Replication of results from young subjects in independent cohort

Using the data from an independent replication cohort of young, healthy subjects (Schott et al., 2014; Assmann et al., 2020), we performed the analyses reported in the main manuscript (see Sections 3 and 4) to infer on optimal modeling of the experimental paradigm and subsequent memory effect. Performing these analyses, we were largely able to replicate results, sometimes with remarkable overlap between original and replication cohort (see Figure S5), sometimes with even stronger evidence for the most often selected model (see Figure S8). Results for the replication cohort are displayed in Figures S5-S10 in this supplement which are designed analogously to Figures S3 and 3-7 in the main paper.

### Supplementary Tables

| **Scanning parameters** | | **FADE**<br>*original cohort* | **yFADE**<br>*replication cohort* |
|---|---|---|---|
| N | number of subjects | 106 young + 153 older | 117 young |
| n | number of scans | 206 | 205 |
| TR | repetition time | 2.58 s | 2.4 s |
| T | total scanning time | 531.48 s | 492 s |
| t | number of trials | 132 | 132 |
| ISI | inter-stimulus-intervals | 0.7 – 2.65 s | 0.5 – 2.3 s |
| | motion correction | realignment & unwarping | realignment only |
| | spatial resolution (native) | 3 x 3 x 3 mm | 2 x 2 x 3 mm |
| | normalized voxel size | 3 x 3 x 3 mm | 3 x 3 x 3 mm |
| | smoothing kernel (FHWM) | 6 mm | 6 mm |

**Table S1.** *Scanning parameters in original and replication cohort.* Overview of fMRI acquisition and data processing parameters in the original cohort ("FADE") and the replication cohort ("yFADE").

| Subjects | Items | 1 | 2 | 3 | 4 | 5 | Hits/ FAs | cHR |
|---|---|---|---|---|---|---|---|---|
| young (N = 106) | old | 10.80 ±8.45 | 14.23 ±9.44 | 11.17 ±8.67 | 13.22 ±8.07 | 50.59 ±16.74 | 63.81 ±14.05 | 52.42 ±14.47 |
| | new | 49.51 ±19.42 | 25.73 ±14.17 | 25.73 ±14.17 | 7.65 ±5.63 | 3.73 ±4.22 | 11.39 ±6.67 | |
| older (N = 153) | old | 15.59 ±10.90 | 6.86 ±7.21 | 10.70 ±8.87 | 8.57 ±8.56 | 58.27 ±16.76 | 66.84 ±13.42 | 45.71 ±13.96 |
| | new | 51.90 ±22.95 | 12.85 ±13.46 | 14.11 ±12.19 | 7.78 ±9.64 | 13.35 ±10.45 | 21.14 ±12.90 | |
| replication (N = 117) | old | 11.15 ±8.75 | 15.00 ±8.06 | 11.70 ±7.54 | 15.37 ±7.67 | 46.79 ±15.71 | 62.15 ±12.53 | 49.93 ±13.95 |
| | new | 46.04 ±18.51 | 26.48 ±14.09 | 15.27 ±10.57 | 8.55 ±6.67 | 3.67 ±4.09 | 12.22 ±8.17 | |
| young vs. older | old | $t = -3.98$, $p < 0.001$ | $t = 6.78$, $p < 0.001$ | $t = 0.42$, $p = 0.673$ | $t = 4.44$, $p < 0.001$ | $t = -3.63$, $p < 0.001$ | $t = -1.74$, $p = 0.083$ | $t = 3.73$, $p < 0.001$ |
| | new | $t = -0.91$, $p = 0.366$ | $t = 7.34$, $p < 0.001$ | $t = -0.53$, $p = 0.600$ | $t = -0.14$, $p = 0.892$ | $t = -10.25$, $p < 0.001$ | $t = -7.94$, $p < 0.001$ | |

**Table S2.** *Behavioral performance in the subsequent memory task.* Average relative response frequencies and their standard deviations across subjects are given in percent (%) for each response option (1-5), together with the hit rate (hits; the proportion of old items rated as 4 or 5), false alarm rate (FAs; the proportion of new items rated as 4 or 5) and corrected hit rate (cHR; hits minus FAs) for young and older subjects from the original cohort as well as young subjects from the replication cohort (see Table S1). The last row lists results from two-sample t-tests (two-tailed, assuming unequal variances) of young against older subjects. Positive t-values indicate significantly larger values for young subjects and negative t-values indicate significantly larger values for older subjects. All p-values significant at the $\alpha = 0.05$ level remained significant when applying the Holm-Bonferroni correction for multiple comparisons (Holm, 1979).

#### Supplementary Figures

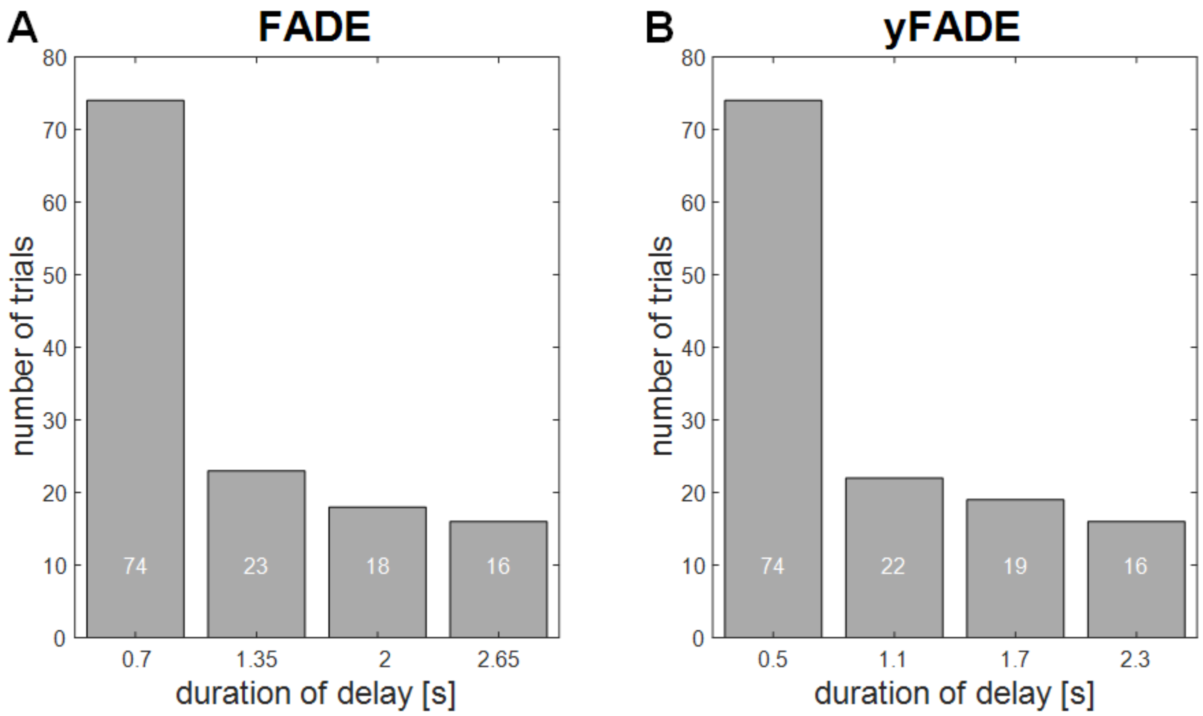


**Figure S1.** *Inter-stimulus-intervals in original and replication cohort.* **(A)** In the original cohort ("FADE"), ISIs were pseudo-exponentially distributed between 0.7 and 2.65 s. This panel is equivalent to Figure 1C from the main paper. **(B)** In the replication cohort ("yFADE"), the distribution of ISIs was almost identical, but bounded between 0.5 and 2.3 s, such that ISIs were a bit shorter on average.

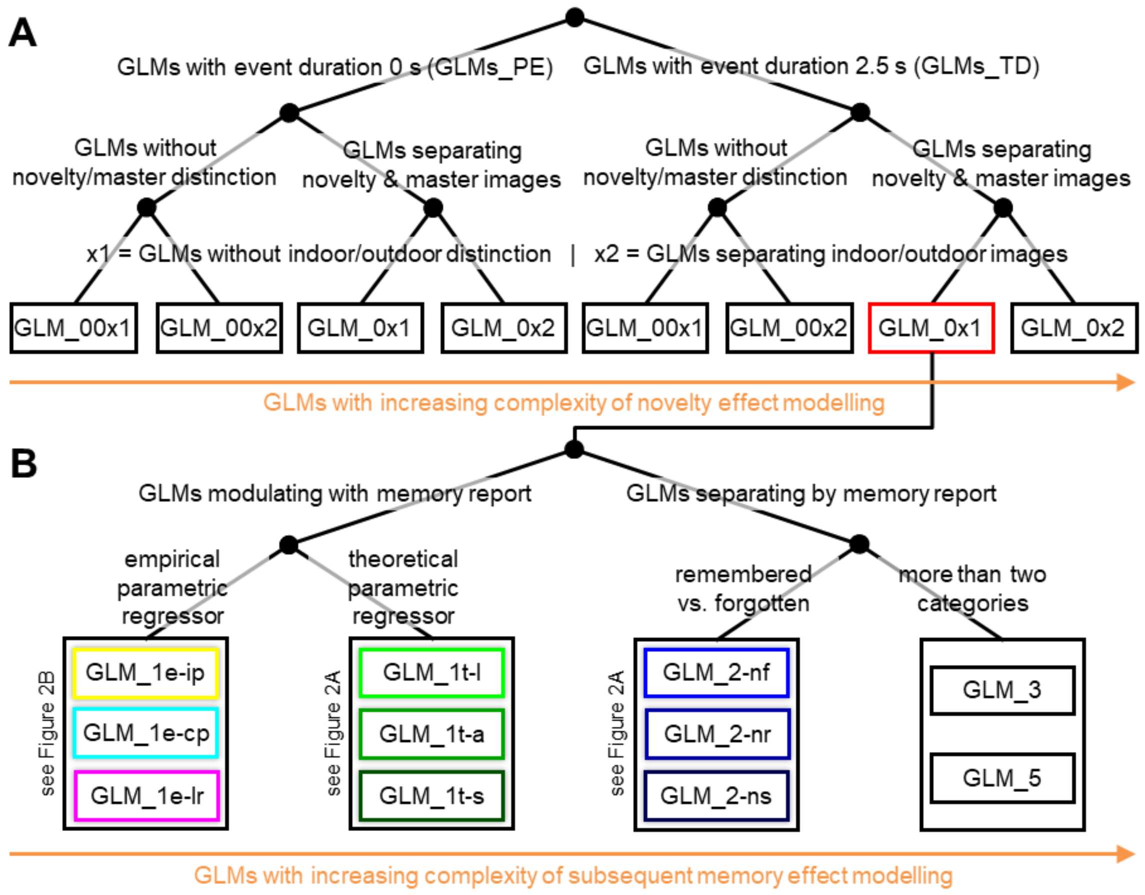


**Figure S2.** *Model space for GLM-based fMRI analyses.* **(A)** 8 models without memory effects varying model features of no interest, namely modeled event duration (top), consideration of stimulus novelty (middle) and consideration of stimulus type (bottom). **(B)** 11 models varying by the way how memory effects are modeled. Each box represents a single first-level GLM; box coloring corresponds to colors used in Figure 2; the box with red outline represents the model referred to as "baseline GLM" in Section 3. For a tabular description of the model space, see Table 1 in the main manuscript.

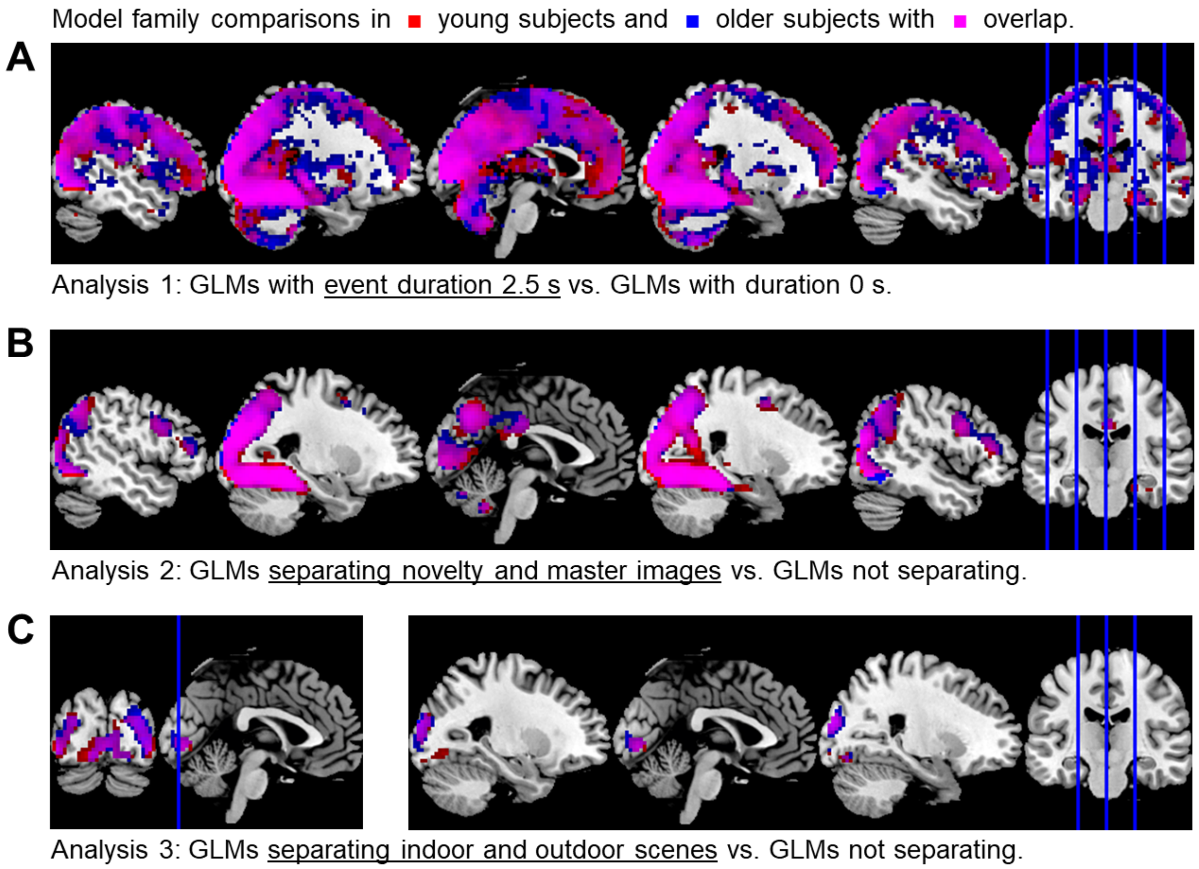


**Figure S3.** *Effects of event duration, novelty and stimulus type.* **(A)** Selected-model maps in favor of GLMs with stimulus length as event duration over point events. **(B)** Selected-model maps in favor of GLMs including a novelty effect. **(C)** Selected-model maps in favor of GLMs assuming an indoor/outdoor effect. Voxels displayed show the respective model preferences in young subjects (red) or older subjects (blue) or both groups (magenta). Selected-model maps display model frequencies and color intensities range from 0 to 1.

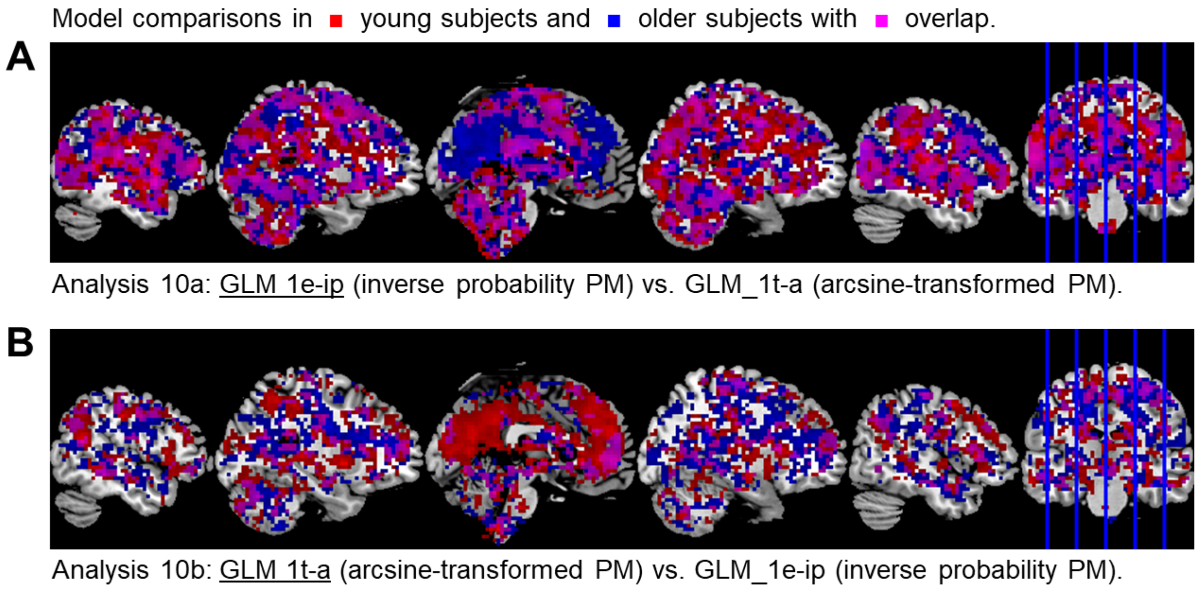


**Figure S4.** *Selected-model maps from winning model comparison.* Selected-model maps in favor of **(A)** the GLM with inverse probability PM and **(B)** the GLM with arcsine-transformed PM, when comparing both against each other. Voxels displayed show the respective model preference in young subjects (red) or older subjects (blue) or both groups (magenta). Selected-model maps display model frequencies and color intensities are ranging from 0 to 1.

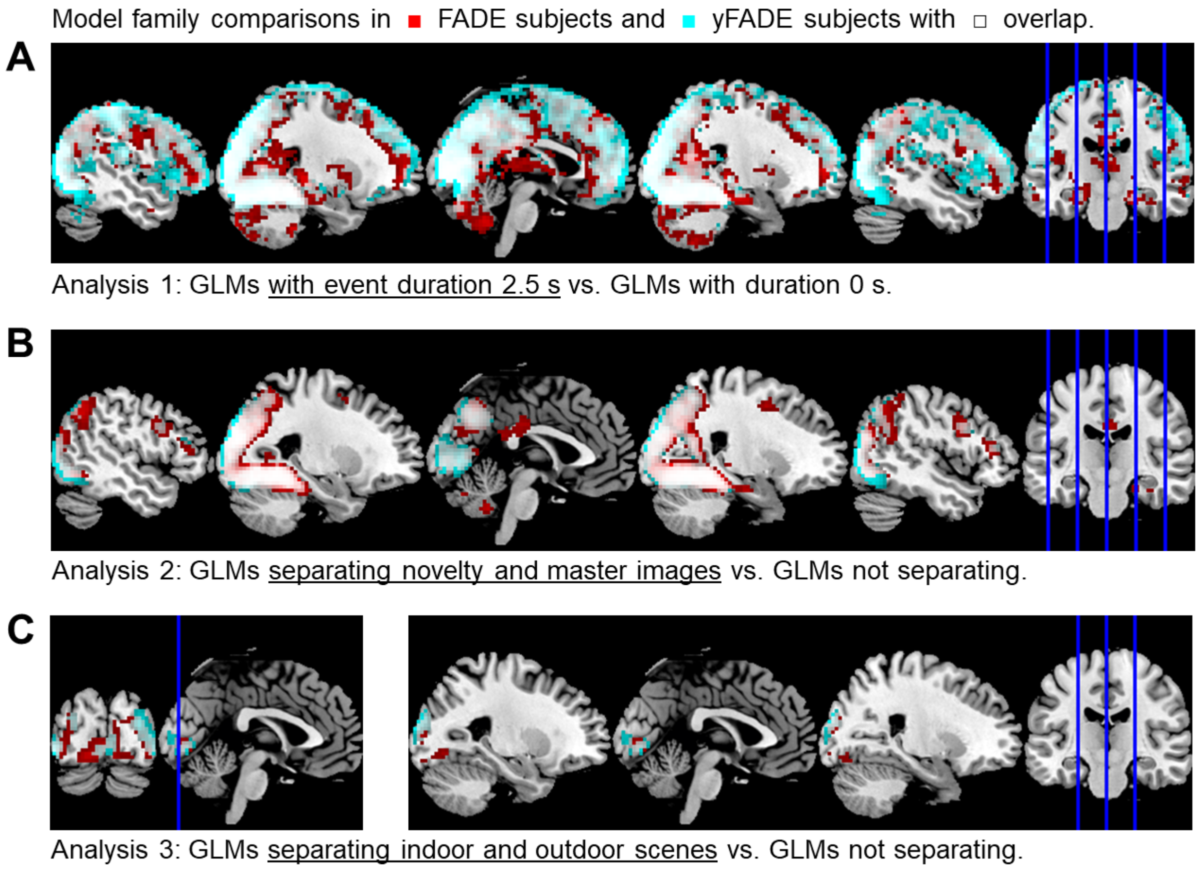


**Figure S5.** *Effects of event duration, novelty and stimulus type in the replication cohort.* **(A)** Selected-model maps in favor of GLMs with stimulus length as event duration over point events. **(B)** Selected-model maps in favor of GLMs including a novelty effect. **(C)** Selected-model maps in favor of GLMs assuming an indoor/outdoor effect. Voxels displayed show the respective model preferences in the original (red), the replication (cyan) or both cohorts (magenta). Selected-model maps display model frequencies and color intensities range from 0 to 1.

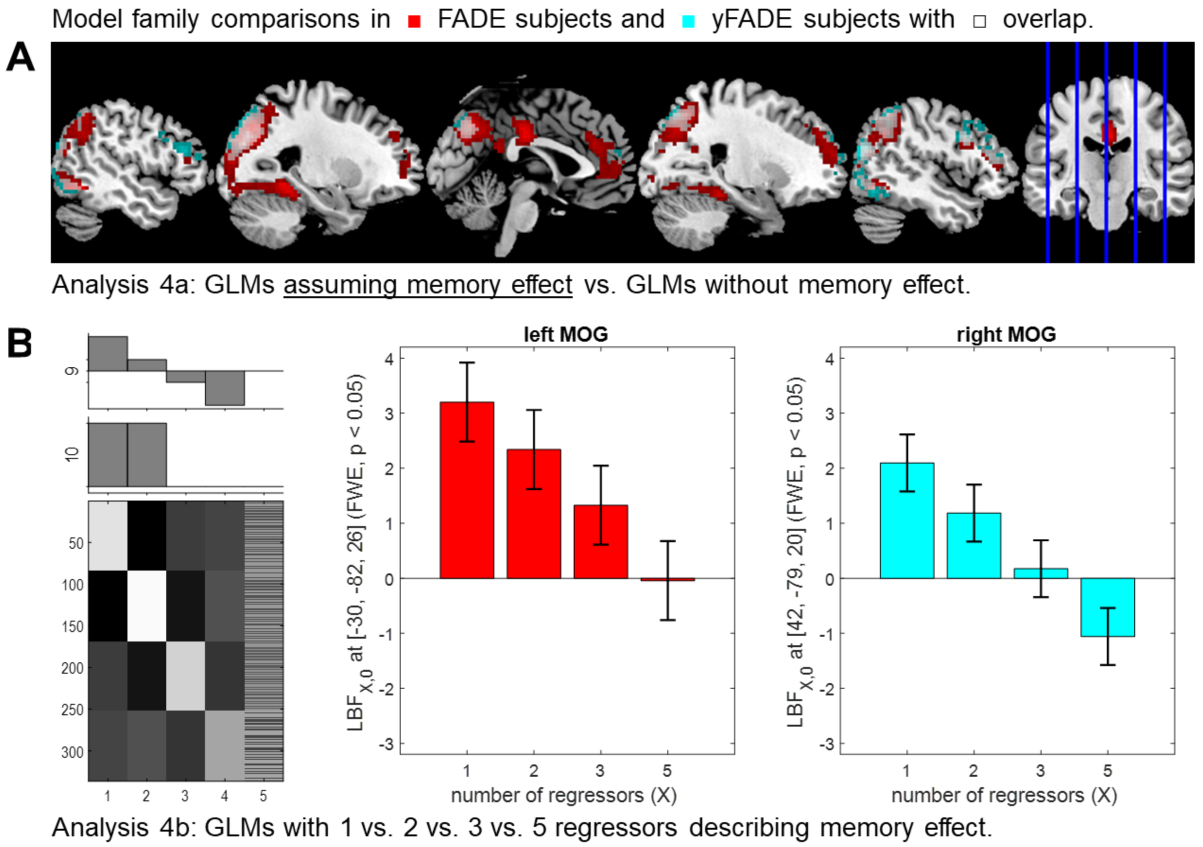


**Figure S6.** *Effects of subsequent memory and number of regressors in the replication cohort.* **(A)** Selected-model maps in favor of GLMs modeling memory using one or two regressors, as obtained from the original (red), the replication (cyan) or both cohorts (magenta). Selected-model maps display model frequencies and color intensities range from 0 to 1. **(B)** Significant linear effects of the number of regressors used to describe memory (X) on the log Bayes factor (LBF) comparing models with X regressors against the baseline GLM, obtained in the global maxima of the respective conjunction contrasts, i.e. the middle occipital gyrus (MOG). Bar plots depict contrasts of parameter estimates of the group-level model; error bars denote 90% confidence intervals (computed using SPM12).

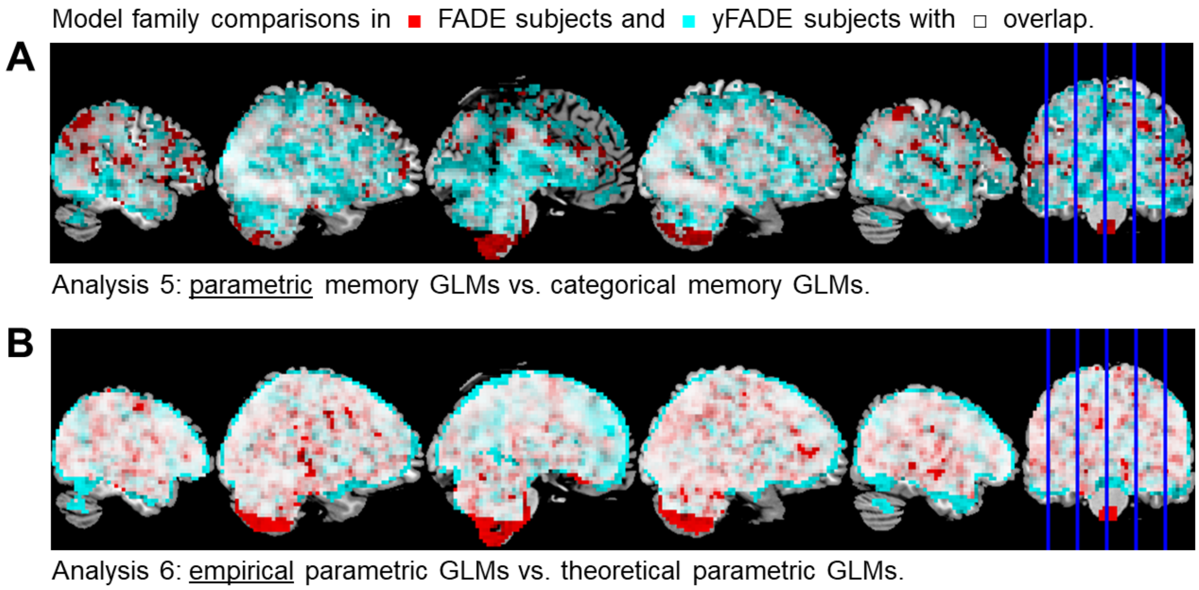


**Figure S7.** *Parametric vs. categorical models of the subsequent memory effect in the replication cohort.* **(A)** Selected-model maps in favor of parametric GLMs against categorical GLMs. **(B)** Selected-model maps in favor of empirical parametric GLMs against theoretical parametric GLMs. Voxels displayed show the respective model preferences in the original (red), the replication (cyan) or both cohorts (magenta).

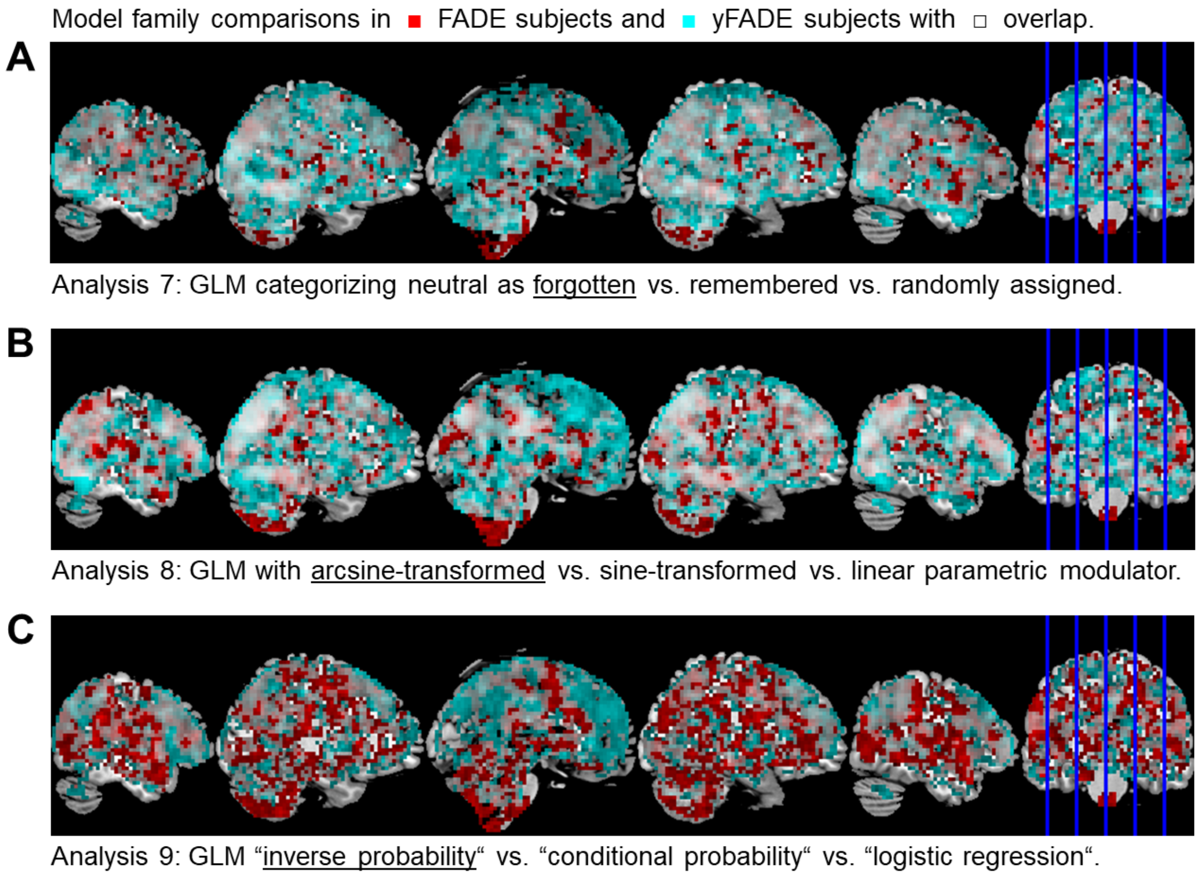


**Figure S8.** *Winning models within model families in the replication cohort.* **(A)** Selected-model maps in favor of the GLM treating neutral images as forgotten items within the two-regressor categorical GLMs. **(B)** Selected-model maps favoring the GLM using an arcsine-transformed parametric modulator within the theoretical parametric GLMs. **(C)** Selected-model maps in favor of the GLM using an inverse probability parametric modulator within the empirical parametric GLMs. Voxels displayed show the respective model preferences in the original (red), the replication (cyan) or both cohorts (magenta).

**Figure S9 (see next page).** *Model (family) comparisons (summary) in the replication cohort.* **(A)** and **(B)** Selected-model maps (original cohort) in favor of GLMs assuming a novelty effect (red; see Figure S3B), a memory effect (blue; see Figure S6A), parametric vs. categorical memory effects (green; see Figure S7A) or an arcsine-shaped subsequent memory effect vs. other theoretical models (magenta; see Figure S8B). In most voxels with preference for parametric GLMs, there was also a preference for the arcsine model. **(C)** and **(D)** The corresponding selected-model maps from the replication cohort. **(E)** Proportion of voxels in which a model or family was selected (original cohort). “X within Y” is to be read as “probability that X was the selected family among voxels in which Y was the selected family”. **(F)** Same proportions as in E, obtained from the replication cohort.

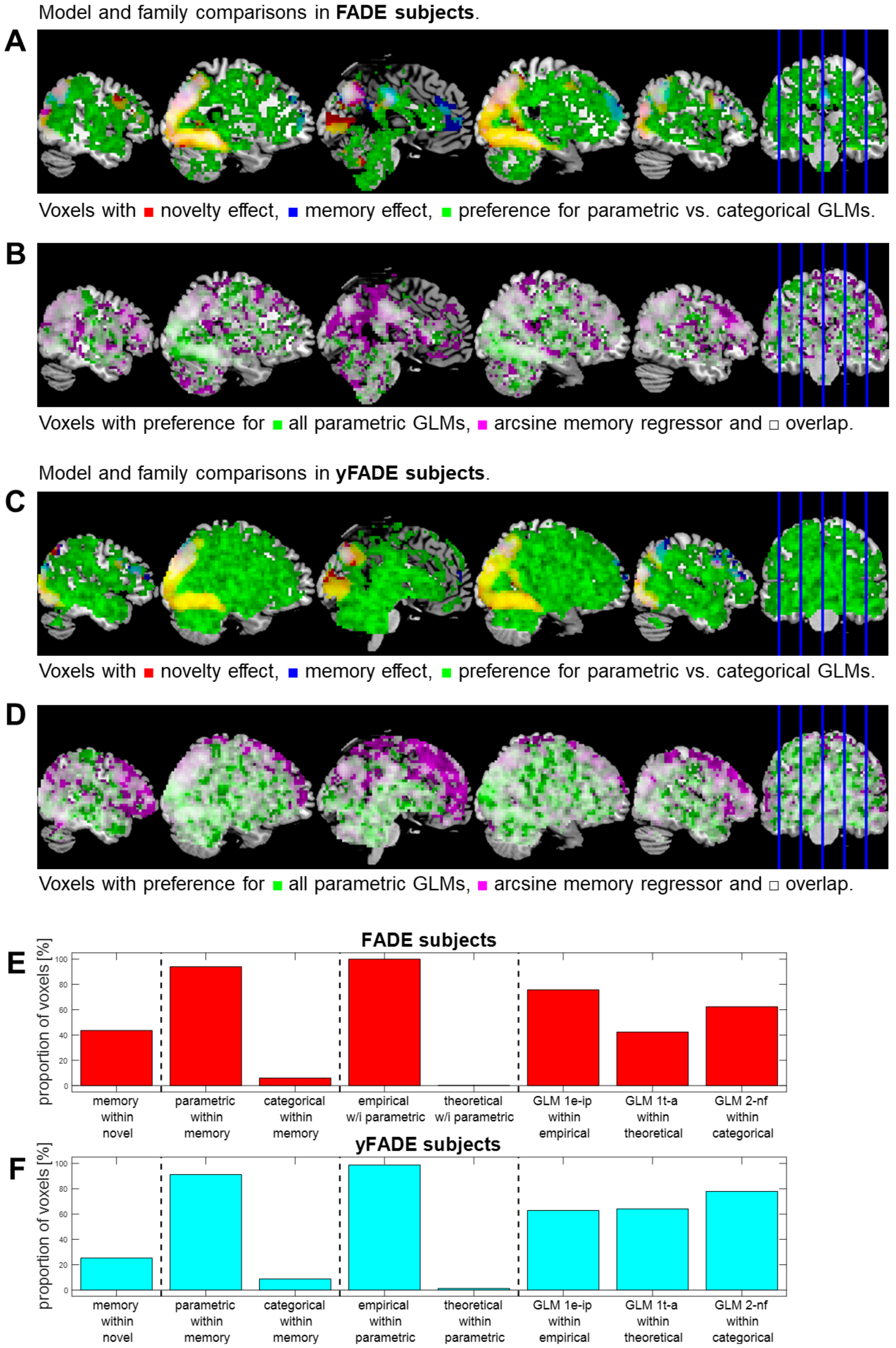
Model and family comparisons in FADE subjects.
A
Voxels with ■ novelty effect, ■ memory effect, ■ preference for parametric vs. categorical GLMs.
B
Voxels with preference for ■ all parametric GLMs, ■ arcsine memory regressor and □ overlap.
Model and family comparisons in yFADE subjects.
C
Voxels with ■ novelty effect, ■ memory effect, ■ preference for parametric vs. categorical GLMs.
D
Voxels with preference for ■ all parametric GLMs, ■ arcsine memory regressor and □ overlap.
E
FADE subjects
proportion of voxels [%]
memory within novel
parametric within memory
categorical within memory
empirical w/i parametric
theoretical w/i parametric
GLM 1e-ip within empirical
GLM 1t-a within theoretical
GLM 2-nf within categorical
F
yFADE subjects
proportion of voxels [%]
memory within novel
parametric within memory
categorical within memory
empirical within parametric
theoretical within parametric
GLM 1e-ip within empirical
GLM 1t-a within theoretical
GLM 2-nf within categorical

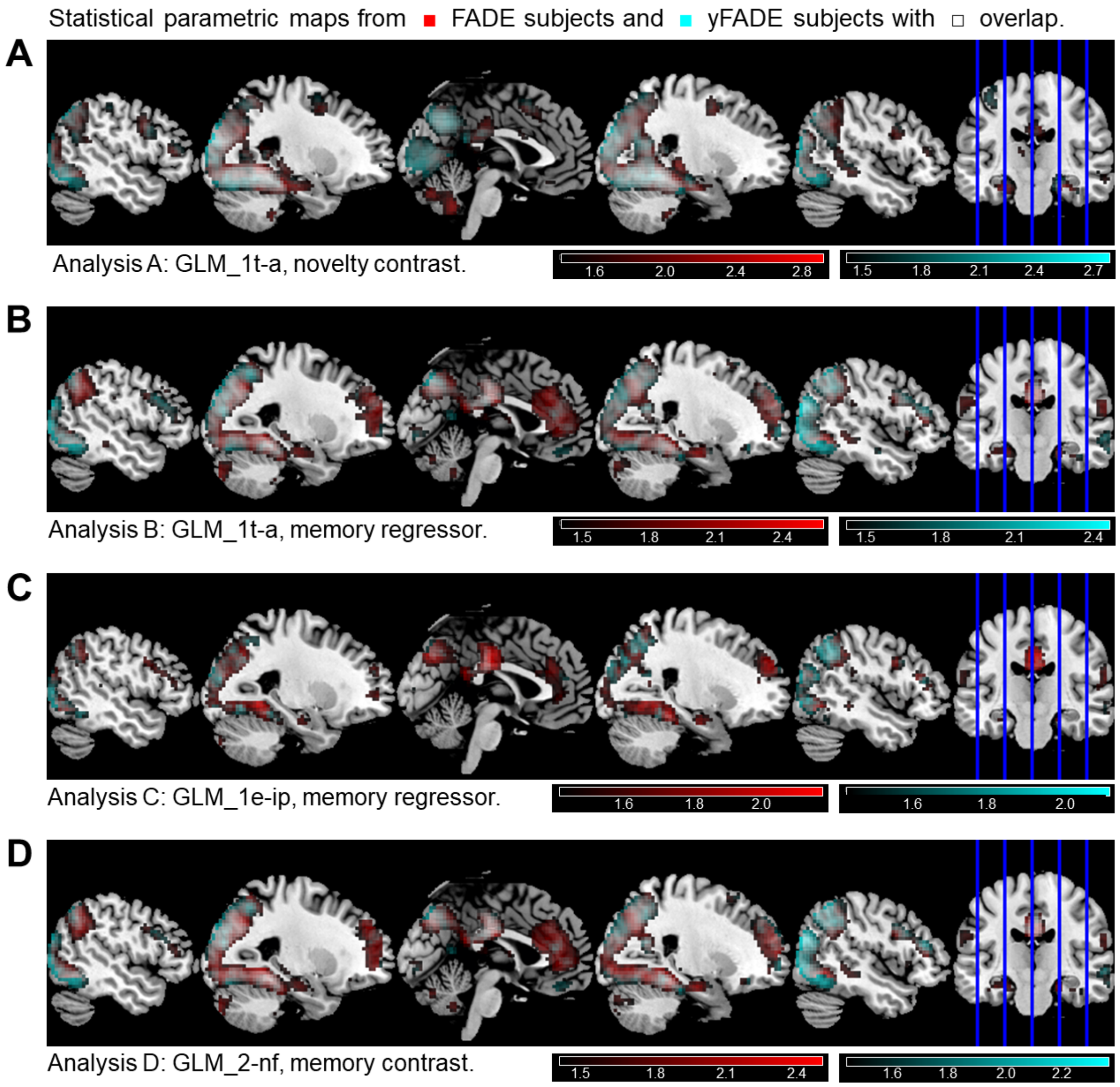


**Figure S10.** *Exemplary statistical parametric maps in the replication cohort.* On the second level, a one-sample t-test was run across parameter estimates obtained from the original cohort (red) and the replication cohort (cyan) for **(A)** the novelty contrast (novelty vs. master images) and **(B)** the memory regressor of the theoretical-parametric GLM using the arcsine-transformed PM, **(C)** the memory regressor of the empirical-parametric GLM using the inverse probability PM and **(D)** the memory contrast (remembered vs. forgotten items) resulting from a two-regressor categorical GLM categorizing neutral responses as forgotten. In SPM, statistical inference was corrected for multiple comparisons (FWE, $p < 0.05$, $k = 10$), resulting in critical F-values for thresholding of SPMs (original: $F > 27.01$; replication: $F > 26.85$). Color maps are scaled from the critical F-value to the maximum F-value in each map, in units of the decadic logarithm (see color bars).

#### References


Assmann A, Richter A, Schütze H, Soch J, Barman A, Behnisch G, Knopf L, Raschick M, Schult A, Wüstenberg T, Behr J, Düzel E, Seidenbecher CI, Schott BH. 2020. Neurocan genome-wide psychiatric risk variant affects explicit memory performance and hippocampal function in healthy humans. *Eur J Neurosci* ejn.14872. doi:10.1111/ejn.14872

Blainey P, Krzywinski M, Altman N. 2014. Replication. *Nat Methods* **11**:879–880. doi:10.1038/nmeth.3091

Eklund A, Nichols TE, Knutsson H. 2016. Cluster failure: Why fMRI inferences for spatial extent have inflated false-positive rates. *Proc Natal Acad Sci U S A* **113**:7900–7905.

Faul F, Erdfelder E, Lang A, Buchner A. 2007. G*Power 3: A flexible statistical power analysis program for the social, behavioral, and biomedical sciences. *Behavior Research Methods* **39**:175-191. doi:10.3758/bf03193146

Holm S. 1979. A simple sequentially rejective multiple test procedure. *Scandinavian Journal of Statistics* **6***:*65-70.

Schott BH, Assmann A, Schmierer P, Soch J, Erk S, Garbusow M, Mohnke S, Pöhland L, Romanczuk-Seiferth N, Barman A, Wüstenberg T, Haddad L, Grimm O, Witt S, Richter S, Klein M, Schütze H, Mühleisen TW, Cichon S, Rietschel M, Noethen MM, Tost H, Gundelfinger ED, Düzel E, Heinz A, Meyer-Lindenberg A, Seidenbecher CI, Walter H. 2014. Epistatic interaction of genetic depression risk variants in the human subgenual cingulate cortex during memory encoding. *Transl Psychiatry* **4**:e372–e372. doi:10.1038/tp.2014.10
